# The injury-induced circular RNA circGLIS3 activates dermal fibroblasts to promote wound healing

**DOI:** 10.1101/2022.09.05.506337

**Authors:** Maria A. Toma, Qizhang Wang, Dongqing Li, Yunting Xiao, Guanglin Niu, Jennifer Geara, Manika Vij, Minna Piipponen, Zhuang Liu, Letian Zhang, Xiaowei Bian, Aoxue Wang, Pehr Sommar, Ning Xu Landén

**Affiliations:** Dermatology and Venereology Division, Department of Medicine Solna, Center for Molecular Medicine, Karolinska Institutet; Stockholm, Sweden; Department of Oromaxillofacial Head and Neck Oncology, Shanghai Ninth People’s Hospital, College of Stomatology, Shanghai Jiao Tong University School of Medicine; Shanghai, China; State Key Laboratory of Oral Diseases, National Clinical Research Center for Oral Diseases, Department of Oral and Maxillofacial Surgery, West China Hospital of Stomatology, Sichuan University; Chengdu, China; Key Laboratory of Basic and Translational Research on Immune-Mediated Skin Diseases, Chinese Academy of Medical Sciences; Jiangsu Key Laboratory of Molecular Biology for Skin Diseases and STIs, Institute of Dermatology, Chinese Academy of Medical Sciences and Peking Union Medical College, Nanjing, China; Department of Dermatology and Venereology, Medical Center – University of Freiburg; Freiburg, Germany; Department of Dermatology, The Second Hospital of Dalian Medical University, College of Integrative Medicine, Dalian Medical University; Dalian, China; Department of Plastic and Reconstructive Surgery, Karolinska University Hospital, Stockholm, Sweden; Ming Wai Lau Centre for Reparative Medicine, Stockholm Node, Karolinska Institutet, Stockholm, Sweden

## Abstract

Delayed skin wound healing and excessive scarring are consequences of an impaired healing process and represent a major health and economic burden worldwide. Current intervention strategies lack efficacy and suffer from high recurrence rates necessitating the investigation into alternative treatment modalities like circular RNAs (circRNAs). By RNA sequencing, we profiled circRNA expression changes during human skin wound healing as well as in keratinocytes and fibroblasts isolated from donor-matched skin and acute wounds. CircGLIS3 was found to be transiently upregulated in the dermal fibroblasts upon skin injury, which was at least partially due to the activated IL-1 signaling. Similarly, overabundant circGLIS3 expression was detected in human keloid lesions compared to the surrounding healthy skin. We found that circGLIS3 resided mainly in the cytoplasm, where it interacted with and stabilized Procollagen C-endopeptidase enhancer 1 (PCPE-1) protein to enhance TGF-β signaling, fibroblast activation, and production of extracellular matrix – important biological processes required for wound repair. Accordingly, knockdown of circGLIS3 in human *ex vivo* wounds potently reduced wound contraction and delayed re-epithelialization. Collectively, we have identified a previously uncharacterized circRNA regulator of human skin wound healing that may open an avenue for circRNA-based therapeutics for abnormal scarring or nonhealing wounds.

**One Sentence Summary:** Transient increase of the circular RNA circGLIS3 promotes the wound fibroblast activation and extracellular matrix production to facilitate wound closure.

## INTRODUCTION

Delayed skin repair and skin fibrosis affect millions of people around the world annually, representing a heavy medical and economic burden (*1*). Despite the high prevalence and the use of different therapeutic approaches for skin repair impairments, no treatments effectively revert or prevent chronic wounds or excessive scars (*2, 3*). Thus, it is critical to elucidate the molecular factors driving healthy skin repair to understand better what mediates its complications.

Skin wound repair is a multiphase process that requires detailed coordination of multiple cell types (e.g., immune, epithelial, stromal) and signaling pathways to achieve healing. During repair, skin cells are subjected to the sequential but overlapping phases of inflammation, growth, and remodeling (*4*). Keratinocytes are the main cellular component of the epidermal layer in the skin, while fibroblasts are the main cell type found in the mostly acellular dermis. Both keratinocytes and fibroblasts produce extracellular matrix (ECM), which is crucial for all the healing phases (*5*).

Circular RNAs (circRNAs) are covalently closed single-stranded RNA molecules that have the 3’ and 5’ ends joined together through back-splicing (*6*). Due to their unique structure, they are more stable than linear RNAs and were shown to have tissue- and cell-specific expression patterns (*7*). Mechanistically, circRNAs sequester microRNAs, interact with proteins, and even encode short peptides (*8, 9*). In the past decade, several studies have begun to explore the functional roles of circRNAs in tissue homeostasis and disease (*10-12*), but too few have addressed, so far, the role of circRNAs in skin repair.

To fill up this knowledge gap and further identify circRNAs with potential functions in wound repair, we performed RNA sequencing (RNA-seq) to profile circRNA expression change during human skin wound healing (*13*) as well as in keratinocytes and fibroblasts isolated from donor-matched skin and acute wounds. We identified circGLIS3 as being transiently upregulated in the wound dermis and overexpressed in skin fibrotic disease keloid, which suggested its potential role in wound fibroblasts. Our following functional study revealed that circGLIS3 promotes fibroblast activation and ECM production by increasing the cellular responsiveness to TGF-β1 signaling. Importantly, we found that human *ex vivo* wounds lacking circGLIS3 failed to close, which further reinforces its essential role in wound repair.

## RESULTS

### CircGLIS3 is upregulated in wound fibroblasts

To identify circRNAs with functional roles in skin repair, we first aimed to profile their expression change during human skin wound healing. For this, we developed a unique human *in vivo* wound model by making full-thickness excisional wounds (3mm in diameter) on the skin of healthy volunteers and then collecting wound-edge tissues with 6mm biopsy punches 1, 7, and 30 days later from the same donor (**Fig. 1A, Table 1**, and **Table S1**). These time points are chosen to capture the three sequential phases of wound repair, i.e., inflammation (∼3 days), proliferation (∼4-21 days), and remodeling (∼21days-one year) (*4*). To further dissect the circRNA expression in individual cell types, we also isolated CD45^-^ epidermal keratinocytes and CD90^+^ dermal fibroblasts from some of these tissue samples by magnetic activation cell sorting (**Fig. 1A**).

**Table 1.** Characteristics of tissue donors (n=40)

| Experiments | Study<br>population | Age, years<br>(mean ± s.d.) | Ethnicity; Sex<br>(male: female) |
| --- | --- | --- | --- |
| Full-thickness biopsy RNA-seq (*Fig. 1B-C*) | 5 | 65.3 ± 3.2 | Caucasian; 2:3 |
| Isolated cell type RNA-seq (*Fig. 1B, D*) | 5 | 33.2 ± 10.3 | Caucasian; 3:2 |
| Full-thickness biopsy qRT-PCR validation<br>(*Fig. 1E*) | 5 | 63.6 ± 3.4 | Caucasian; 0:5 |
| Isolated cell type qRT-PCR validation (*Fig. 1G-H*) | 5 | 33.8 ± 12.1 | Caucasian; 3:2 |
| LCM-qRT-PCR validation (*Fig. 1F*) | 7 | 40.6 ± 13.3 | Caucasian; 4:3 |
| Skin for *ex vivo* wound model (*Fig. 6*) | 3 | 49 ± 6.6 | Caucasian; 0:3 |
| Skin for fibroblast isolation (*Fig. 2-5*) | 2 | 50 ± 1 | Caucasian; 0:2 |
| Skin and keloid biopsies from patients (*Fig. 1I*) | 8 | 36,8 ± 11,9 | Asian; 4:4 |

840 Abbreviation: s.d., standard deviation

**Fig. 1.**
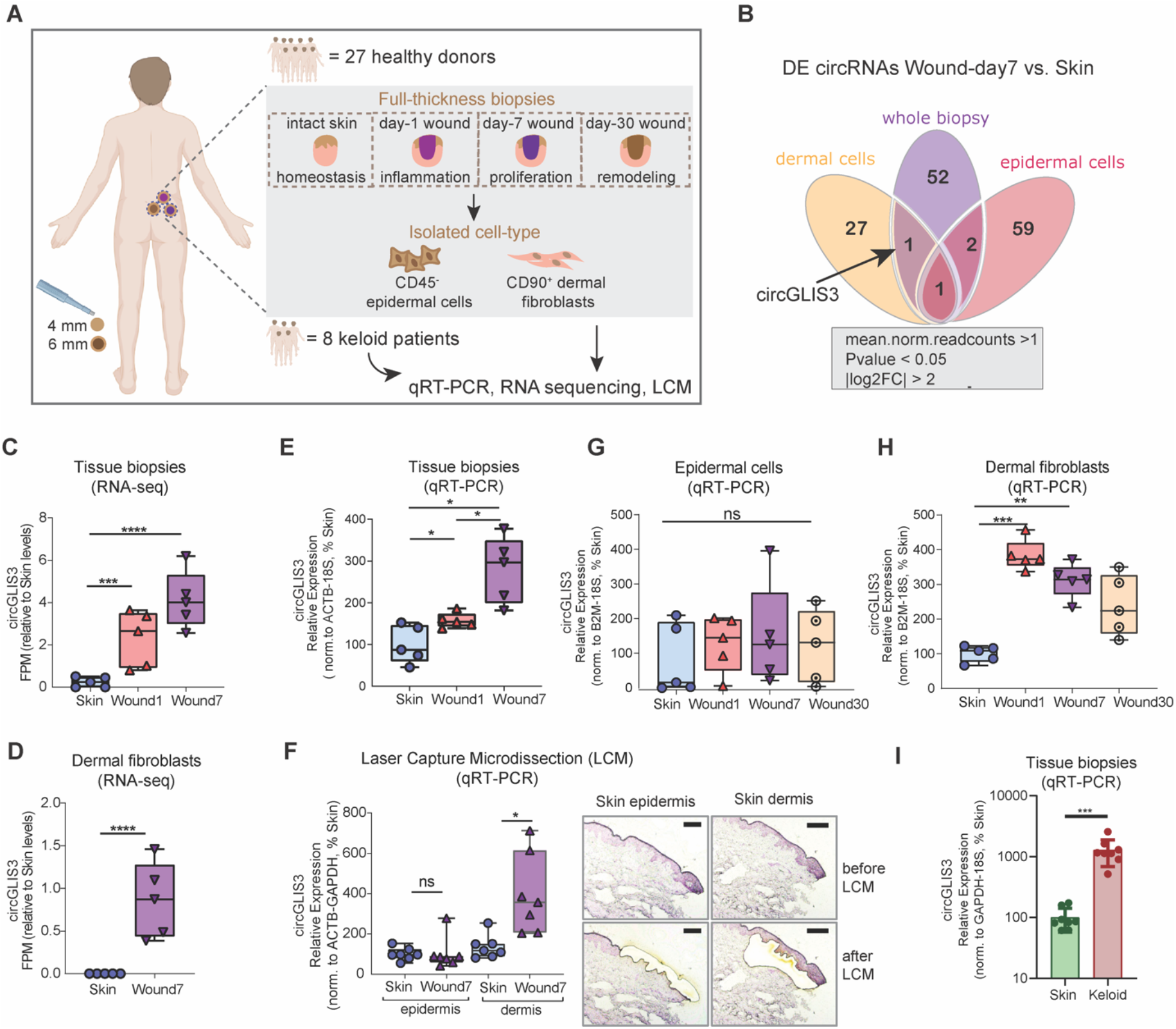
CircGLIS3 is upregulated in wound fibroblasts. **(A)** Excisional wounds were created on the skin of 27 healthy volunteers and collected 1 (Wound1), 7 (Wound7), and 30 days later (Wound30) from the same donor. CD45^-^ epidermal cells and CD90^+^ fibroblasts were isolated from matched skin and Wound7 samples. Biopsies were also collected from lesional sites and surrounding skin of 8 keloid patients. CircRNAs were analyzed in these clinical samples by RNA-seq, qRT-PCR, and laser capture microdissection (LCM). **(B)** Venn diagram showing the commonly identified differentially expressed (DE) circRNAs in the isolated cell types and tissue biopsies of the skin and Wound7 analyzed by RNA-seq. CircGLIS3 expression in the skin and wound tissue biopsies (n=5 donors) **(C)** and isolated fibroblasts (n=5 donors) **(D)** was analyzed by RNA-seq. qRT-PCR validation of circGLIS3 expression in additional skin and wound biopsies (n=5 donors) **(E)**, LCM of epidermal and dermal compartments of the skin and wounds (n=7 donors) **(F)**, CD45^-^ epidermal cells **(G)** and CD90^+^ fibroblasts **(H)** isolated from the skin and wounds (n=5 donors), donor matched skin and keloid biopsies (n=8 donors) **(I)**. *P<0.05, **P<0.01, ***P<0.001, and ****P<0.0001 by Wilcoxon test **(D, I)** or RM one-way ANOVA and Tukey’s multiple comparisons test **(C, E, F**-**H)**.

We profiled circRNA expression by poly (A) independent total RNA-seq in the matched skin and day7-wound tissues (n=5 donors) as well as the isolated cells (n=5 donors). By filtering the top changed and abundant circRNAs (normalized read counts >1, *P* < 0.05 by Wald test, and |log2FoldChange| > 2), we identified 56 differentially expressed (DE) circRNAs in the day7-wound compared to the skin tissues, 62 and 29 DE circRNAs in keratinocytes and fibroblasts, respectively, isolated from the day7-wound versus skin (**Fig 1B** and **Table S2**). By intersecting the sequencing results of the tissues and cells, we found circGLIS3 to be the only circRNA upregulated in wound biopsies and fibroblasts compared to the skin and whose expression was not altered in wound keratinocytes (**Fig. 1B-D** and **Table S2**).

We further validated circGLIS3 expression in additional clinical samples (**Table 1** and **Table S1)** by quantitative real-time PCR (qRT-PCR) with divergent primers to specifically amplify the back-splicing junction (BSJ) of circGLIS3, which is absent in its cognate linear isoform (**Fig. 2A**). In the wound and skin tissues (n=5 donors) (**Fig. 1E)**, epidermal and dermal compartments separated by laser capture microdissection (LCM) of the skin and wound-edges (n=7 donors) (**Fig. 1F)**, as well as epidermal cells and dermal fibroblasts sorted from the matched skin and wounds at each stage of wound healing (n=5 donors) **(Fig. 1G, H)**, we confirmed the prominently upregulated circGLIS3 expression in dermal fibroblasts, but not in epidermal cells, during skin wound healing.

**Fig. 2.**
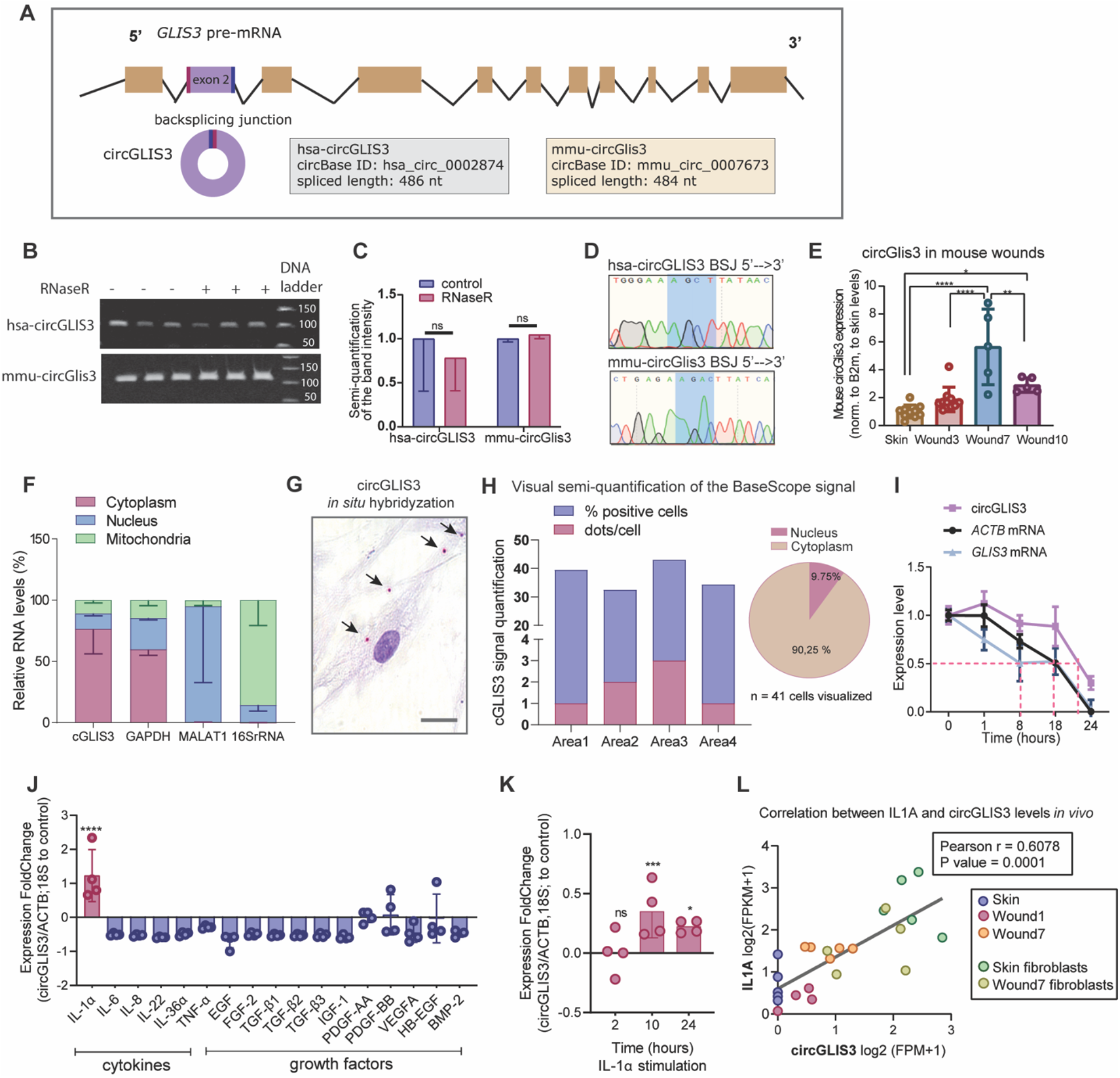
The molecular characteristics of circGLIS3. (**A**) Illustration of circGLIS3 biogenesis. (**B**) Agarose gel electrophoresis of circGLIS3 RT-PCR products from RNaseR-digested or control RNA from human (top) and mice (bottom) fibroblasts (n=3). Band intensity was quantified in (**C**). (**D**) Sanger sequencing of the RT-PCR products verified sequences of the BSJ regions of human (top) and mouse (bottom) circGLIS3. (**E**) qRT-PCR of circGLIS3 in skin and day3, 7, and 10 acute wounds from C57BL/6 mice (n=15). (**F**) qRT-PCR of circGLIS3, *GAPDH, MALAT1*, and *16S* rRNA in nuclear, cytoplasmic, and mitochondrial fractions of human fibroblasts (n=3). (**G**) *In situ* hybridization of circGLIS3 in human fibroblasts. (**H**) Visual semi-quantification of the circGLIS3 positive cells and the number of circGLIS3 signal dots/cell. (**I**) qRT-PCR of circGLIS3, *GLIS3*, and *ACTB* mRNA levels in human fibroblasts treated with Actinomycin-D (n=4). qRT-PCR of circGLIS3 in human fibroblasts treated with wound-related cytokines and growth factors for 24 hours (n=4) (**J**), or IL-1α for 2-24 hours (n=4) (**K**). (**L**) Correlation between circGLIS3 and *IL1A* expression in human wound samples analyzed by RNA-seq. Data are presented as means±SD. *P<0.05, ***P<0.001, and ****P<0.0001 by two-tailed Student’s t-test (**J, K**) or RM one-way ANOVA and Tukey’s multiple comparisons test (**E**).

Interestingly, in keloids, a fibroproliferative skin disease characterized by an abnormal wound healing process, massive production of ECM, and hyperplasia of dermal tissue (*14, 15*), we also found overabundant circGLIS3 expression in lesional sites compared to donor-matched healthy skin biopsies (n=8 donors) (**Fig. 1I, Table 1**, and **Table S1)**. These results highlight the specific upregulation of circGLIS3 expression in wound fibroblasts and skin fibrosis, which prompted us to further study the role of circGLIS3 in dermal fibroblasts.

### Molecular characterization of circGLIS3

CircGLIS3 (circBase ID: hsa_circ_0002874) is a circular RNA that derives from the second exon of the *GLIS family zinc finger 3 (GLIS3)* gene, and its expression is conserved in other species such as mouse (**Fig. 2A**) (*16*). We confirmed its circularity by using primers spanning the BSJ of either human or mouse circGLIS3 in RT-PCR analysis (**Fig. 2B** and **C, Table S4**). The circularity of circGLIS3 was further supported by its resistance to the exonuclease RNase R digestion, which in turn leads to the degradation of most linear RNAs (**Fig. 2B** and **C**) (*17*). The sequences of predicted BSJ regions of human and mouse circGLIS3 were verified by Sanger sequencing of the RT-PCR products (**Fig. 2D**). Moreover, in a mouse *in vivo* wound model, we found that circGlis3 expression was also upregulated in the day-7 and day-10 acute wounds compared to the intact skin (**Fig. 2E**). Thus, the injury-induced circGLIS3 expression pattern is conserved between human and mouse.

We further characterized the subcellular localization of circGLIS3 to elucidate its mode of action. For this, we divided human primary fibroblasts into subcellular fractions of nucleus, cytoplasm, and mitochondria, which were enriched with nuclear long non-coding RNA MALAT1 (*18*), *GAPDH* mRNA, and *16S* mitochondrial rRNA (*19*), respectively, confirming a successful fraction separation. We found that circGLIS3 was mainly detected in the cytoplasm (**Fig. 2F**), which was also observed in *in situ* hybridization analysis of circGLIS3 in fibroblasts (**Fig. 2G, H**, and **Fig. S1A**).

Moreover, we treated fibroblasts with Actinomycin-D (5µg/mL) to block transcription (*20*) and then characterized the stability of circGLIS3 (**Fig. 2I**). qRT-PCR analysis revealed a longer half-life of 22 hours for circGLIS3 compared to 8 hours for GLIS3 mRNA or 18 hours for ACTB mRNA (**Fig. 2I**). The above analysis revealed that circGLIS3 was an exonic circRNA, resistant to RNaseR digestion (*21*), more stable compared to linear RNAs (*22*), and localized mainly in the cell cytoplasm.

To understand why circGLIS3 expression was increased upon skin injury, we stimulated human dermal fibroblasts with a panel of cytokines and growth factors that are reportedly important for wound repair (**Fig. 2J**). We found that IL-1α treatment led to circGLIS3 upregulation (**Fig. 2J**), which occurred less than 10 hours after adding IL-1α ( **Fig. 2K**). IL-1 is rapidly produced in the injured site and is one of the first signals to alert the surrounding cells of barrier damage (*23*). Interestingly, qRT-PCR results showed that *GLIS3* mRNA levels were not altered by IL-1α, indicating that IL-1 signaling may specifically affect circGLIS3 biogenesis and does not enhance the expression of the *GLIS3* gene (**Fig. S1B**). Additionally, we found that IL-1α and circGLIS3 expression levels were significantly correlated (Pearson r= 0.6078, P = 0.0001) in human skin and wounds *in vivo* as well as in the isolated fibroblasts from skin and wounds analyzed by RNA-seq (**Fig. 2L**). These results suggest that the increased circGLIS3 expression in wound fibroblasts may be a consequence of the injury-induced IL-1α signal activation.

### CircGLIS3 enhances TGF-β1 signaling

To investigate the role of circGLIS3 in skin wound healing, we modulated its expression in human dermal fibroblasts. To knock down circGLIS3 expression, we designed three siRNAs targeting its diagnostic junction (si-circGLIS3). As controls, a scrambled siRNA sequence (si-ctrl) and a siRNA partially (10 out of 21 nucleotides) complementary to the junction sequence of circGLIS3 (si-circGLIS3_ctrl) were used (**Fig. S2A, Table S3**). To overexpress circGLIS3, we subcloned the DNA sequence corresponding to circGLIS3 and its endogenous flanking region, which also included complementary circular frames needed for circularization, into a plasmid expression cassette (pLC5-circGLIS3). With qRT-PCR analysis, we confirmed that both strategies effectively changed the circGLIS3 levels in fibroblasts and did not significantly affect those of its linear counterparts (**Fig. S2B-E**).

We next carried out transcriptomic profiling by microarray in dermal fibroblasts and identified 56 up- and 74 down-regulated genes (|log2FC| > 1.5, FDR < 0.05) upon circGLIS3 knockdown (**Fig. 3A**). Gene Ontology enrichment analysis of these differentially expressed genes (DEGs) revealed biological processes important for wound healing and placed TGF-β1 among the top signaling pathways affected by circGLIS3 (**Fig. 3B**). TGF-β1 is a crucial growth factor involved in many biological processes essential for wound repair, such as inflammation, angiogenesis, granulation tissue formation, and remodeling (*24*). TGF-β1 is known to potently induce the differentiation of fibroblasts into the more contractile myofibroblasts as well as the deposition and remodeling of ECM (*25*). Gene Set Enrichment Analysis (GSEA) of our microarray data revealed that the genes involved in the TGF-β1 pathway (from GSE79621, a dataset of TGF-β1-induced transcriptional response in human dermal fibroblasts) (*26*) were significantly enriched among the genes downregulated by the circGLIS3 knockdown in fibroblasts, suggesting that circGLIS3 may potentiate TGF-β1 signaling (**Fig. 3C**).

**Fig. 3.**
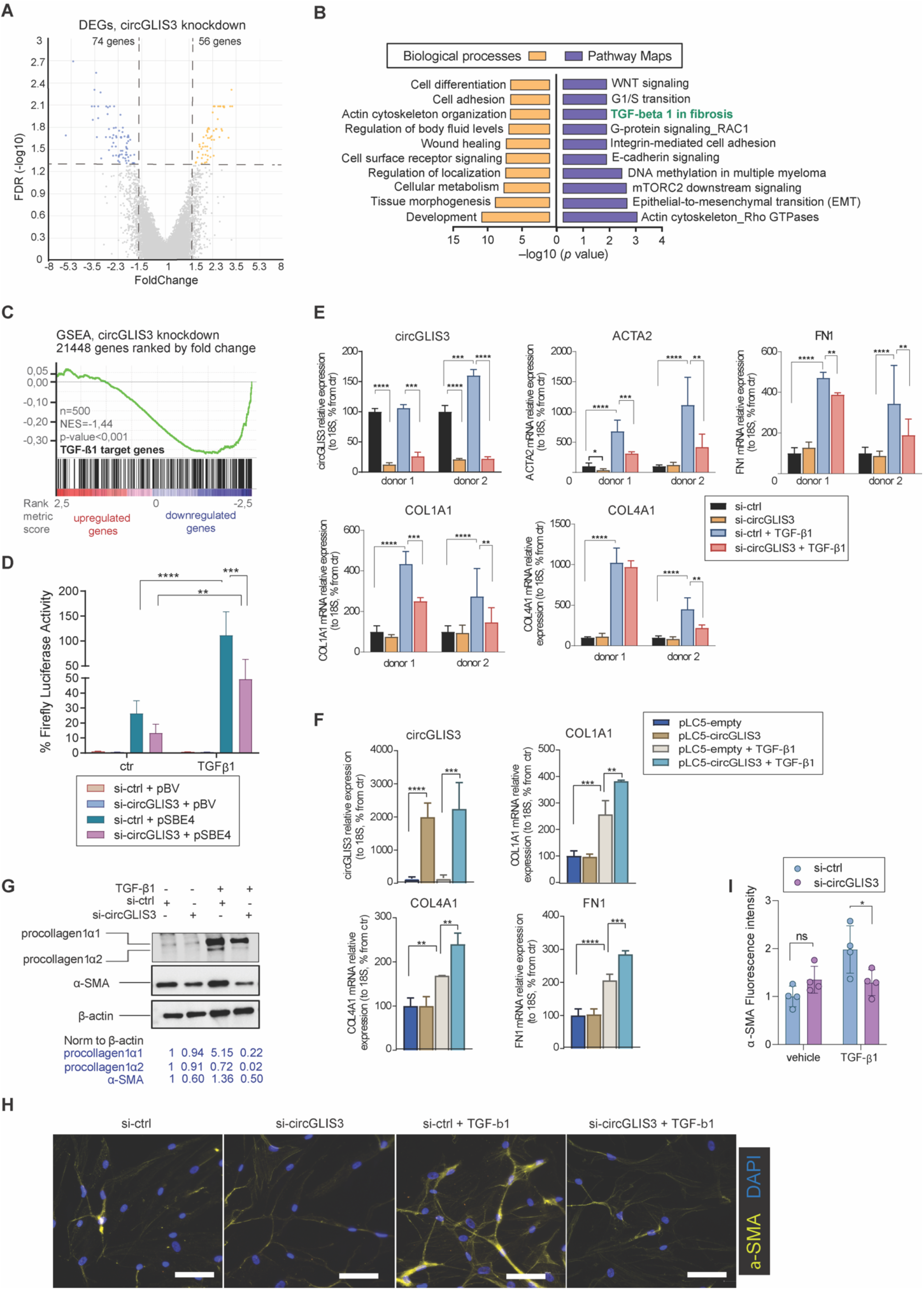
CircGLIS3 enhances TGF-β1 signaling. (**A**) Microarray profiling of human fibroblasts with circGLIS3 knockdown. The volcano plot shows the differentially expressed genes (DEGs) with |Fold change|>1.5 and FDR<0.05. (**B**) Gene Ontology analysis of the DEGs. (**C**) Gene Set Enrichment Analysis evaluated the enrichment of TGF-β1 signaling-related genes in the microarray data. (**D**) Luciferase activity in fibroblasts transfected with TGF-β reporter plasmid or empty vector together with si-ctrl or si-circGLIS3 for 24 hours and then treated with TGF-β1 for another 24 hours (n=4). qRT-PCR of circGLIS3, *ACTA2, FN1, COL1A1*, and *COL4A1* mRNA in fibroblasts transfected with si-ctrl or si-circGLIS3 (**E**), circGLIS3 overexpression plasmid or empty vector (**F**) for 24 hours and then stimulated with TGF-β1 for another 24 hours (n=4). (**G**) Western blotting of procollagen type 1 and α-SMA and semi-quantification of the band intensity (relative to β-actin levels) in fibroblasts with circGLIS3 depletion and TGF-β1 treatment. (**H**) Immunofluorescence staining of α-SMA in fibroblasts with circGLIS3 depletion and TGF-β1 treatment. Scale bar=100 µm. The signal intensity was quantified in (**I**). Data are presented as means±SD (**D-F, I**). **P<0.01, ***P<0.001, and ****P<0.0001 by one-way ANOVA and Dunnett’s multiple comparisons test (**D-F**); *P<0.05 by two-tailed Student’s t-test (**I**).

To assess whether circGLIS3 directly affects the activity of the TGF-β1 pathway, we co-transfected siRNAs targeting circGLIS3 with a luciferase reporter construct containing multiple copies of TGF-β response transcriptional activator Smad binding elements (pSBE4) into human primary fibroblasts (*27*). We showed that TGF-β1 treatment enhanced the expression of this luciferase reporter (**Fig. 3D**). Importantly, circGLIS3 silencing significantly reduced the luciferase activity under the TGF-β1 treatment, demonstrating that circGLIS3 is required for the TGF-β1 signaling in fibroblasts (**Fig. 3D**). Accordingly, we found that the expression of several TGF-β1-induced target genes, including alpha smooth muscle actin (*ACTA2*, also referred to as α-SMA), fibronectin 1 (*FN1*), collagen type I (*COL1A1*) and IV (*COL1A4*), were significantly downregulated by circGLIS3 knockdown (**Fig. 3E**), whereas their expression was further enhanced by circGLIS3 overexpression (**Fig. 3F**). Moreover, we observed that circGLIS3 silencing also decreased the TGF-β1-induced α-SMA and procollagen type 1 protein expression, as shown by Western blotting (**Fig. 3G**) and immunofluorescence (**Fig. 3H** and **I**). Additionally, in murine dermal fibroblasts with circGlis3 silencing, we observed a decrease in the Tgf-β1-induced expression of Acta2, Col1a1, Col4a1, and Fn1, which parallels the effect observed in human fibroblasts (**Fig. S3**). Collectively, our results identified circGLIS3 as a positive regulator for TGF-β1-induced fibroblast activation into matrix-secreting fibroblasts.

### CircGLIS3 interacts with and stabilizes PCPE-1

We next explored the molecular mechanism by which circGLIS3 regulates fibroblast activation. As circGLIS3 was mainly detected in the cytosol (**Fig. 2F-H**), we sought to characterize its protein interactome. Due to limited transfection efficiency in primary fibroblasts, we performed circRNA pulldown in HEK293T cells by expressing circGLIS3 tagged with MS2 hairpins and a FLAG-tagged fusion protein recognizing MS2 (MS2-CP). The circGLIS3-protein complexes were pulled down by FLAG antibody-conjugated beads, and the co-purified proteins were subjected to mass spectrometry (MS) analysis (**Fig. 4A**). The successful pulldown of the MS2-tagged circGLIS3 compared to the non-tagged circGLIS3 was confirmed by qRT-PCR analysis of circGLIS3 (**Fig. 4B**). We identified 55 proteins uniquely bound to the circGLIS3-MS2, and among them, Procollagen-C Proteinase Enhancer 1 (PCPE-1) and Rho GTPase Activating Protein 31 (RHG31) were enriched in dermal fibroblasts, as shown in the Human Protein Atlas (**Fig. 4C, Table S4**, and **S6**). Compared to RHG31, PCPE-1 had a higher interaction score with circGLIS3-MS2 and more abundant expression in fibroblasts (**Fig. 4D** and **Table S5**). In human dermal fibroblasts, the circGLIS3-PCPE-1 interaction was further validated by RNA-binding protein immunoprecipitation (RIP), where we precipitated PCPE-1 protein with an antibody (**Fig. 4E**). We showed that circGLIS3, but not *GAPDH* mRNA, was enriched in the anti-PCPE-1 group compared to the IgG negative control (**Fig. 4F**).

**Fig. 4.**
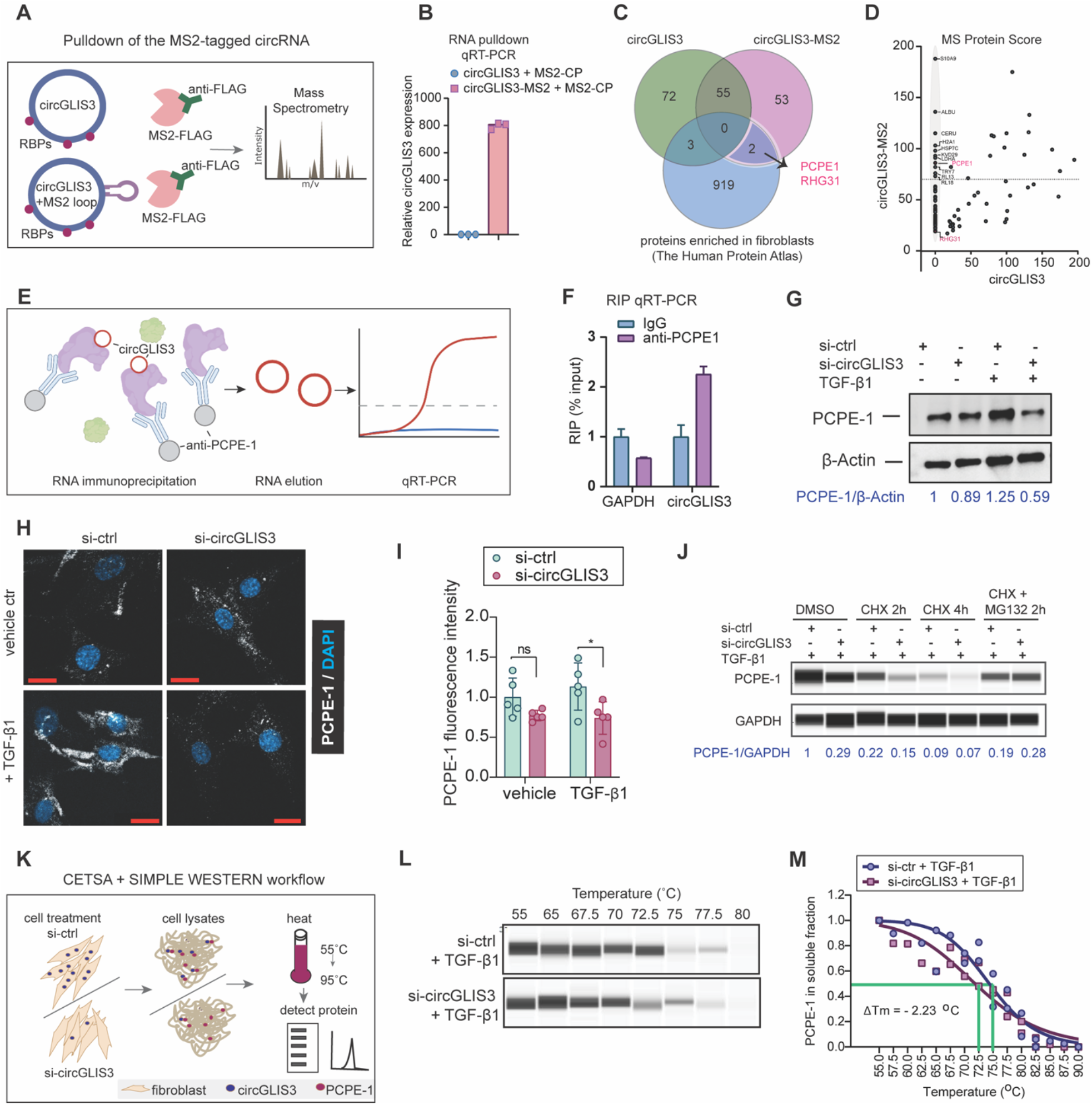
CircGLIS3 interacts with and stabilizes PCPE-1. (**A**) A plasmid containing circGLIS3 tagged with MS2 hairpins (circGLIS3-MS2) was co-transfected with a plasmid expressing a FLAG-tagged fusion protein with an MS2-recognizing portion (MS2-CP) in cells. Control cells were co-transfected with circGLIS3 plasmids and MS2-CP. The ribonucleoprotein complexes (RNPs) were isolated by using anti-FLAG antibodies, and the eluted RNA-binding proteins (RBD) were analyzed by Mass Spectrometry (MS). (**B**) qRT-PCR of circGLIS3 in the RNPs. (**C**) Venn diagram showing proteins identified by MS overlapped with a list of proteins expressed in human dermal fibroblasts. (**D**) Proteins identified by MS were plotted with their interaction scores. (**E**) Schematics of RNA-binding protein immunoprecipitation (RIP) strategy. (**F**) qRT-PCR of circGLIS3 and *GAPDH* mRNA in RNPs immunoprecipitated with anti-PCPE-1 antibody or IgG. Western blotting (**G**) and immunofluorescence analysis (IF) (**H, I**) of PCPE-1 in fibroblasts with circGLIS3 depletion and TGF-β1 treatment. Scale bar=20 µm. (**J**) Simple Western of PCPE-1 in fibroblasts with circGLIS3 depletion and TGF-β1 treatment for 24 hours and then treated with cycloheximide (CHX) and/or MG132 for 2-4 hours. (**K**) Illustration of CETSA assay. Simple Western (**L**) and melting curves (**M**) of PCPE-1 protein in the CETSA assay. *P<0.05 by two-tailed Student’s t-test.

PCPE-1 has been known for its TGF-β1-induced expression and its important function in collagen biosynthesis (*28*). Here, Western blotting (**Fig. 4G**) and immunofluorescent staining (**Fig. 4H** and **I**) showed that PCPE-1 protein levels were reduced by circGLIS3 knockdown in fibroblasts treated with TGF-β1. Thereby, we hypothesized that circGLIS3 might be needed to maintain the level of PCPE-1 proteins by either enhancing its production or reducing its degradation. To discern these two possibilities, we probed the endogenous PCPE-1 protein turnover in TGF-β1-stimulated fibroblasts treated with an inhibitor of protein translation – cycloheximide (CHX), and a proteasome inhibitor – MG132 (*29*). We found that blockage of the protein degradation pathway, but not inhibition of translation, equalized the PCPE-1 protein levels between circGLIS3-depleted fibroblasts and control fibroblasts, suggesting the importance of circGLIS3 for stabilizing the PCPE-1 protein (**Fig. 4J**). To validate this, we performed CEllular Thermal Shift Assay (CETSA) to assess whether circGLIS3 may affect the thermal properties of PCPE-1 protein (**Fig. 4K**) (*30*). Protein lysates from TGF-β1-treated fibroblasts with or without circGLIS3 knockdown were incubated at different temperatures (ranging from 55-90°C), and the amount of PCPE-1 present in the soluble fraction was quantified by Simple Western **(Fig. 4K-M)**. We found that circGLIS3 silencing induced a shift to a lower melting temperature of PCPE-1 (ΔTm = −2,23°C), confirming that circGLIS3 is needed to stabilize PCPE-1 protein in fibroblasts (**Fig. 4M**).

Furthermore, we interrogated the role of PCPE-1 on TGF-β1 signaling in dermal fibroblasts. Similar to circGLIS3, knockdown of PCOLCE, the gene encoding the PCPE-1 protein, also decreased the TGF-β1-induced expression of the matrisome genes, such as *COL1A1, COL4A1*, and *FN1*, and the contractility-related gene, *ACTA2* (**Fig. 5A**-**C**). Moreover, with the TGF-β1 signal-responsive luciferase reporter assay, we showed that lack of PCOLCE also decreased luciferase activity in human primary fibroblasts under both basal and TGF-β1-treated conditions (**Fig. 5D**). Together, these results suggest that circGLIS3 interacts with and stabilizes PCPE-1 protein, which is required for enhancing the TGF-β1 signaling in human dermal fibroblasts (**Fig. 7**).

**Fig. 5.**
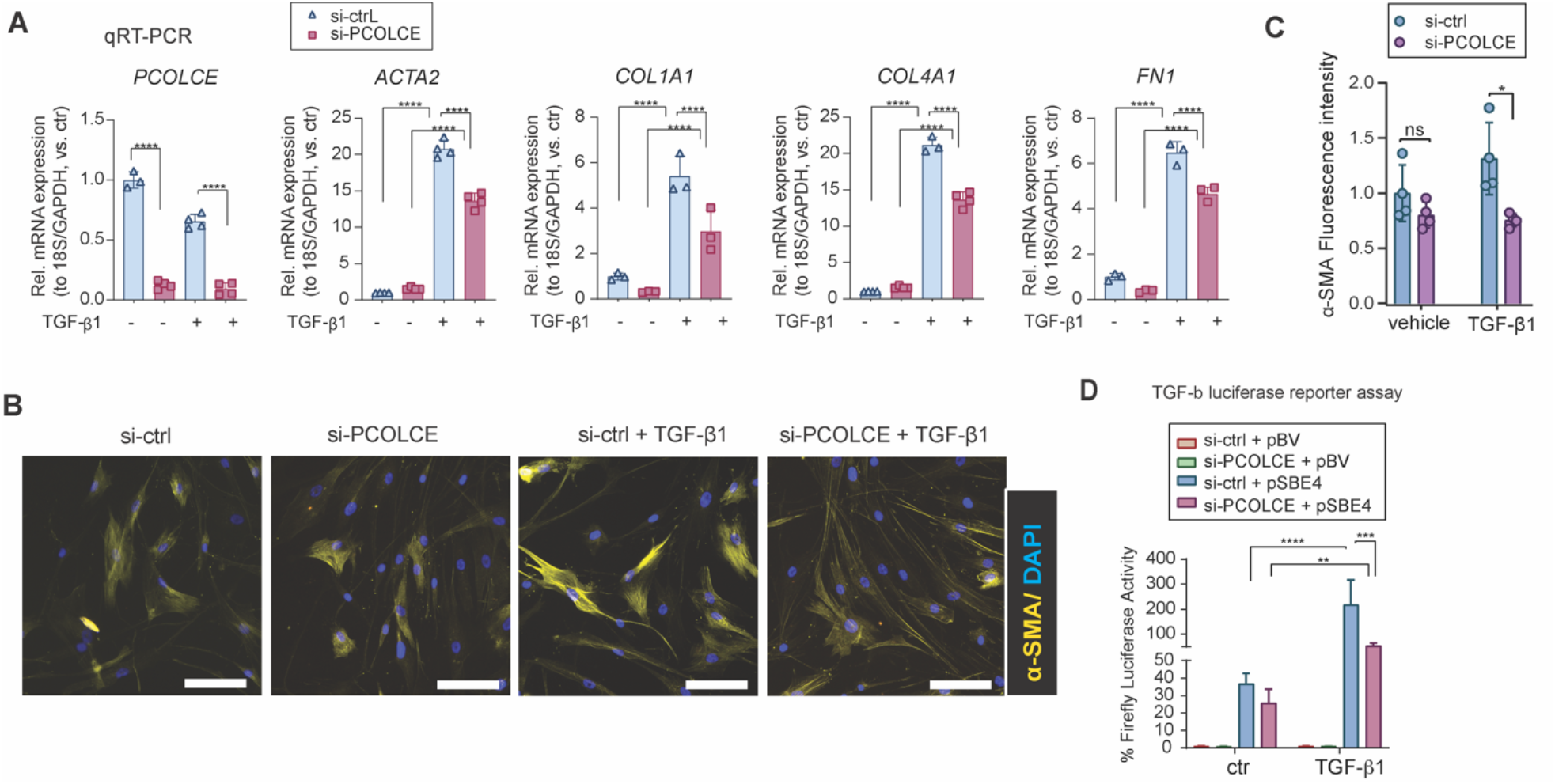
PCPE-1 is required for TGF-β1 signaling and fibroblast activation. **(A)** qRT-PCR of *PCOLCE* (the gene encoding PCPE-1), *ACTA2, COL1A1, COL4A1*, and *FN1* mRNA in fibroblasts transfected with si-ctrl or si-PCOLCE and stimulated with TGF-β1 for 24 hours (n=3-4). Immunofluorescence analysis of α-SMA in human fibroblasts with PCPE-1 depletion and TGF-β1 treatment. Representative pictures (Scale bar = 100 µm) are shown in (**B**), and the signal intensity is quantified in (**C**). (**D**) Luciferase activity in fibroblasts transfected with a TGF-β reporter plasmid or empty vector and si-ctrl or si-PCOLCE for 24 hours and then treated with TGF-β1 (5 ng/µΑ) for another 24 hours. **P < 0.01, ***P < 0.001, and ****P < 0.0001 by one-way ANOVA and Dunnett’s multiple comparisons test.

### CircGLIS3 is needed for wound closure in human *ex vivo* wounds

To further evaluate the potential importance of circGLIS3 in human skin wound healing, we employed a human *ex vivo* wound model (**Fig. 6A**). To this end, we topically applied circGLIS3-specific siRNAs or scramble control oligos on partial-thickness wounds created on surgery discarded human skin immediately after the injury and three days later. On day 6, these wounds were collected for histological and molecular analysis (**Fig. 6A**). We found that the circGLIS3 siRNA treatment hindered wound closure by significantly reducing the re-epithelialization and wound-edge contraction, as shown by hematoxylin and eosin (H&E) staining (**Fig. 6B, C**). Moreover, by qRT-PCR analysis on the separated dermal and epidermal layers of the wound tissues, we confirmed that circGLIS3 levels were effectively reduced by the siRNA treatment in the dermis but not in the epidermis (**Fig 6D, E**). In line with our *in vitro* findings (**Fig. 3E, H**), circGLIS3 knockdown decreased the expression of the contractility-related gene *ACTA2* and the matrisome genes *COL1A1* and *COL4A1* in human *ex vivo* wound dermis (**Fig. 6E**). Additionally, immunofluorescent staining showed reduced α-SMA expression in the dermal compartments of the human *ex vivo* wounds lacking circGLIS3, which explained why these wounds contracted less (**Fig. 6F, G**). Collectively, this study emphasizes the essential role of circGLIS3 in human skin wound healing to promote fibroblast activation and their differentiation into matrix-secreting and contracting cells.

**Fig. 6.**
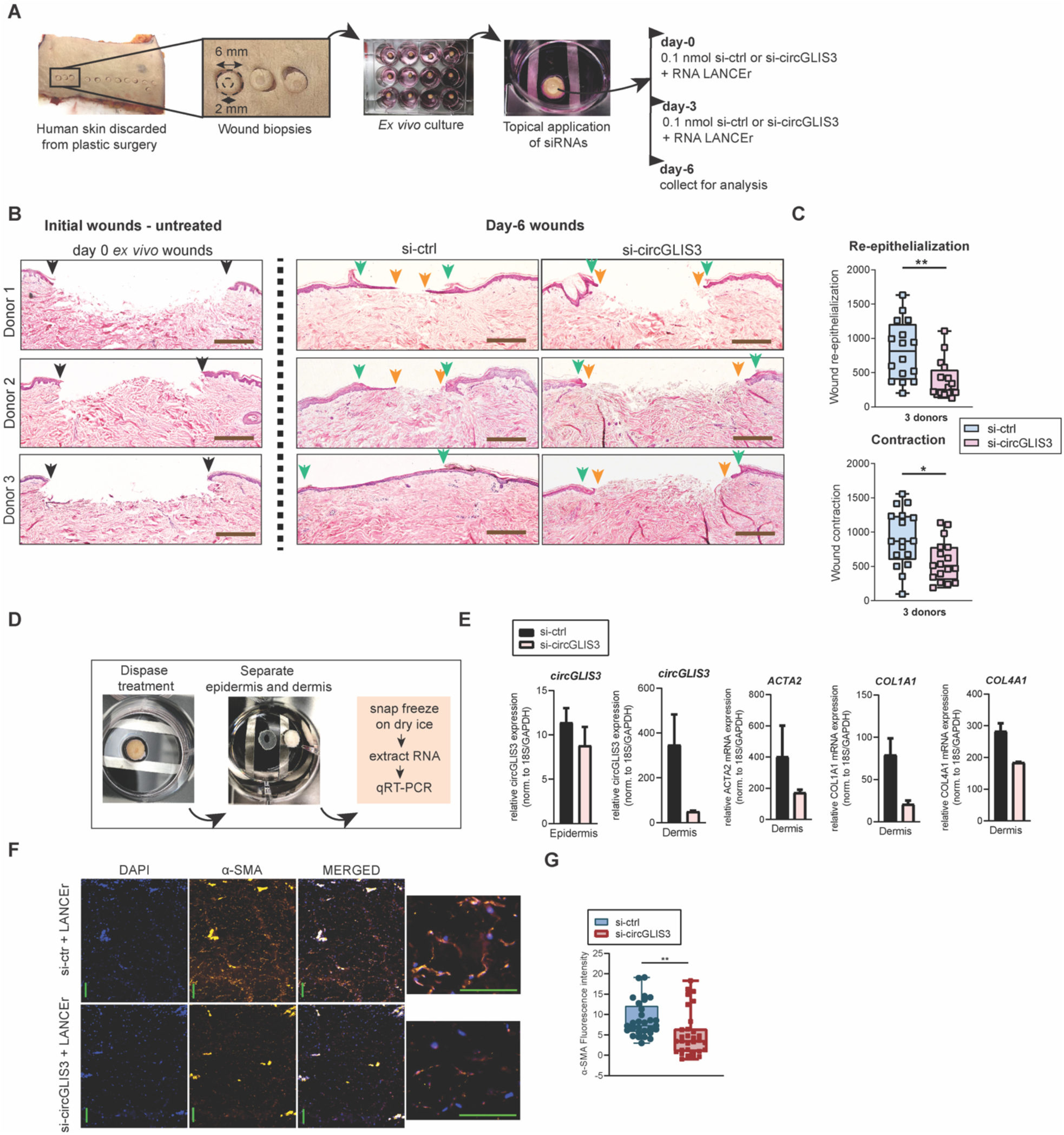
CircGLIS3 is needed for the closure of human *ex vivo* wounds. (**A**) Scheme of topical treatment of human *ex vivo* wounds with si-RNA targeting circGLIS3 or a scrambled control siRNA. (**B**) Hematoxylin and eosin staining of day-0 and day-6 wounds. Black arrows demarcate the initial wound edges at day 0, green arrows indicate the wound edges at day 6, and orange arrows highlight the newly formed epidermis. Scale bar = 500 μm. (**C**) Quantification of wound re-epithelialization (distance between green arrows – distance between orange arrows) and contraction (distance between black arrows – distance between green arrows) of at least two wounds per donor for three donors. (**D**) Workflow of epidermis and dermis separation from day-1 *ex vivo* wounds. (**E**) qRT-PCR of circGLIS3 expression in the epidermis and dermis and *ACTA2, COL1A1*, and *COL4A1* mRNA in the dermis of *ex vivo* wounds (n = 2 donors). Immunofluorescence analysis of α-SMA on the treated *ex vivo* wound. Representative pictures (scale bar = 100 µm) are shown in (**F**), and the signal intensity is quantified in (**G**). *P < 0.05, **P < 0.01 by two-tailed Student’s t-test (**C, G**).

**Fig. 7.**
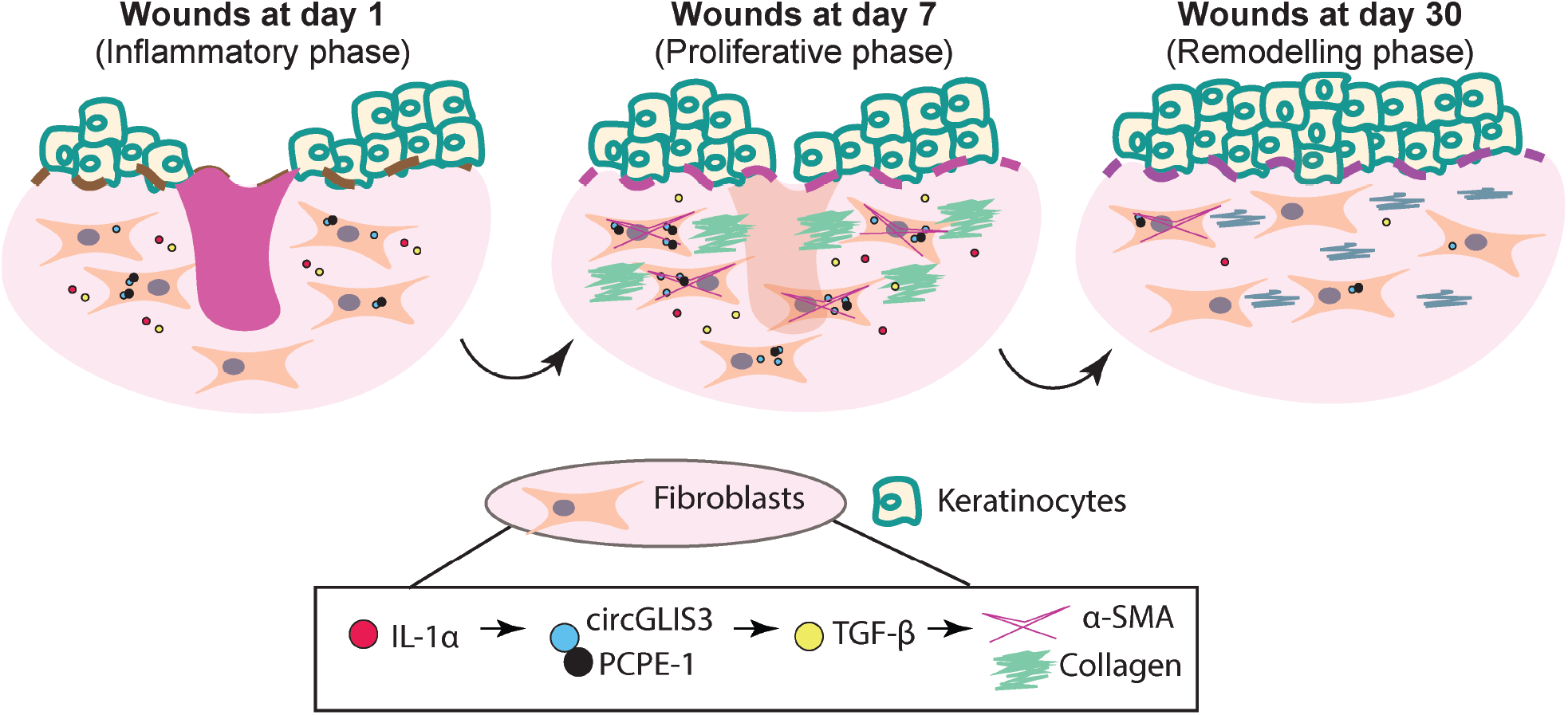
Summary of the study findings. circGLIS3 expression is transiently upregulated in the dermal fibroblasts upon skin injury, which was at least partially due to the activated IL-1 signaling. circGLIS3 resides mainly in the cytoplasm, where it interacts with and stabilizes PCPE-1 protein, enhancing TGF-β signaling, fibroblast activation, and extracellular matrix production. At the remodeling phase, circGLIS3 expression decreases and approaches the skin fibroblast levels, reinforcing its important role in the early stages of wound healing, where it modulates wound contraction and ECM deposition.

## DISCUSSION

This study provides evidence for the dynamic regulation of circGLIS3 in fibroblasts during skin wound healing, whose expression is upregulated in an IL-1α-dependent manner during early phases and later approaches homeostatic skin levels by day 30. The increased expression of circGLIS3 in the wounds of both humans and mice suggests its evolutionary conserved role in skin repair. Loss- and gain-of-function studies of circGLIS3 demonstrated its role in regulating the cellular responsiveness to TGF-β1, leading to the activation of fibroblasts and the production of ECM– requirements for granulation tissue formation and skin repair (**Fig. 7**) (*31*).

TGF-β1 is released in the wound environment by platelets, macrophages, and keratinocytes at the early stages of healing to promote the recruitment of inflammatory cells and angiogenesis (*23*). In the interim stages of wound healing and the transition to the proliferative phase, TGF-β1 prompts the expression of crucial ECM proteins, such as fibronectin and collagens, and enhances fibroblast contraction via the expression of α-SMA, to enable wound closure (*24, 25, 32*). Despite its beneficial role for wound healing, overexuberant granulation tissue function due to persistent TGF-β1 signaling was shown to lead to excessive scarring (*33, 34*). Keloid scars have been widely associated with exacerbated ECM deposition characterized by excessive fibril collagen and fibronectin, while hypertrophic scars (HTS) also displayed an overwhelming presence of α-SMA-expressing (myo)fibroblasts (*14, 15*). Interestingly, here we identified circGLIS3 to be highly upregulated in keloid lesions compared to the healthy skin found in proximity. Similarly, HTS tissues were shown to contain higher circGLIS3 levels compared to healthy skin tissue (*35*). These lines of evidence highlight that, besides its beneficial role for wound closure, circGLIS3 may also regulate dermal fibrosis and represent a therapeutic target in keloids and HTS.

CircGLIS3 has been recently shown to play oncogenic roles in non-small cell lung cancer, bladder cancer, and glioma (*36-39*). Also, it is packaged into beta cell-derived exosomes and transferred to islet endothelial cells, reducing angiogenesis and contributing to type 2 diabetes development (*40*). Here we identified circGLIS3 as a critical factor controlling activation mechanisms of wound fibroblasts. Mechanistically, it binds to PCPE-1 protein to amplify ECM production via TGF-β1 signaling. PCPE-1 has been known as a secreted protein that enhances collagen maturation by promoting the activity of bone morphogenetic protein 1/tolloid-like proteinases to cleave the C-propeptides from procollagens (*41*). This process is important for the formation of collagen monomers capable of forming fibrils. PCPE-1 has been proposed as a marker of fibrosis, given its consistent overexpression in various fibrotic diseases (*42-44*). In addition, it has been reported that PCPE-1, together with collagen type I and IV and fibronectin, are secreted in higher amounts by HTS fibroblasts compared to normal fibroblasts (*45*), suggesting that PCPE-1 may be a potential therapeutic target. Our findings reveal that the depletion of circGLIS3 in dermal fibroblasts destabilizes PCPE-1 protein, and the reduced PCPE-1 levels compromised TGF-β1 signaling. Thus, the inhibition of circGLIS3 with clinically approved siRNA in dermal fibroblasts may represent a promising therapeutic intervention for reducing scar formation.

IL-1α is a proinflammatory cytokine rapidly released from epidermal keratinocytes at the inflammatory stage of wound healing, and it promotes keratinocyte migration and proliferation (*23*). Additionally, IL-1α has been shown to act in a paracrine fashion to activate dermal fibroblasts and enhance their production of collagen (*46*) and keratinocyte growth factor FGF-7, thus facilitating wound re-epithelialization (*47*). However, despite its beneficial roles for wound closure, increased exposure to IL-1α has been reported to be associated with (*48*) and lead to dermal fibrosis (*49*). Our study uncovers that IL-1α upregulates circGLIS3 expression, which at least partially explains the increased levels of circGLIS3 in fibroblasts in human wounds *in vivo*. Further functional analysis of circGLIS3 suggests that it may mediate some of the pro-healing and pro-fibrotic functions of IL-1α in fibroblasts. Given the high levels of both IL-1α and circGLIS3 in keloids and HTS, we postulate that, while it is beneficial for wound closure, sustained IL-1α/circGLIS3 stimulatory axis in fibroblasts may lead to pathological scarring.

Our study builds upon the emerging roles of circRNAs in skin repair. We have previously shown that a circular RNA deriving from the *PRKDC* locus, hsa_circ_0084443, was upregulated in diabetic foot ulcers (DFUs), which is a common type of chronic nonhealing wounds, compared to acute wounds, and it impaired epidermal keratinocyte migration while promoting their abnormal growth (*50*). Subsequent studies revealed that hsa_circ_0084443 knockdown enhanced keratinocyte migration via miR-31/FBN1 (*51*) and miR-20a-3p/RASA1 (*52*) axes to promote wound healing, reinforcing the therapeutic potential of this circRNA. Another circRNA, circ-Amotl1, was shown to promote fibroblast proliferation and migration and accelerate skin wound healing in mice by facilitating the transcription factor Stat3 nuclear translocation and modulating Dnmt3a and miR-17 function (*53*). Collectively, these studies highlight that circRNAs are potent gene expression regulators required for wound healing. Our current study exposes a previously uncharacterized circRNA player in human skin wound repair and its connection with the crucial TGF-β1 signaling pathway to regulate fibroblast functions.

The human *ex vivo* model employed in this study is clinically relevant; however, it only allows for the study of the early phases of wound healing, not scar formation or fibrosis. Using this model, we uncovered a clear physiological role for circGLIS3 in wound healing, and its importance under pathophysiological conditions, such as keloid, warrants further studies. Previous evidence has shown that circGLIS3 can be exported out of cells (*36, 40*), and it would be of interest to investigate whether circGLIS3 can be found in exosomes alongside PCPE-1 from fibroblasts to mediate collagen maturation and deposition.

In summary, our results comprehensively characterize the function and mechanism of circGLIS3 in dermal fibroblasts, highlighting how the transient upregulation of circGLIS3 is beneficial for skin wound healing. CircGLIS3 induces fibroblast activation via TGF-β1 to increase ECM production and speed up wound closure, which may also contribute to pathological skin scarring. Future work should explore the targeted modulation of circGLIS3 expression to alter ECM production in pathological conditions such as excessive scars.

## MATERIALS AND METHODS

### Study design

The goals of this study were (i) to identify circRNAs with potential functions in human skin wound healing and (ii) to uncover the physiological role of circGLIS3 in wound fibroblasts and its underlying molecular mechanism. For circRNA identification and quantification by RNA-seq, LCM, and qRT-PCR, human skin and wound biopsies were obtained from healthy volunteers, and matched skin and lesion biopsies were obtained from patients with keloids. Written informed consent was obtained from all the donors for collecting and using clinical samples. The study of donors 1-32 was performed at the Karolinska University Hospital Solna (Sweden) and was approved by the Stockholm Regional Ethics Committee. The keloid and matched skin tissue samples from donors 33-40 were obtained from the Jiangsu Biobank of Clinical Resources (China) and approved by the Ethics Committee of the Hospital for Skin Diseases (Institute of Dermatology), Chinese Academy of Medical Sciences and Peking Union Medical College. The study was conducted according to the Declaration of Helsinki’s principles. A series of *in vitro* experiments were performed to assess gene expression, cell function, and RNA-protein interactions on dermal fibroblasts isolated from human skin. A human *ex vivo* wound model was used to study the impact of circGLIS3 on wound healing in an *in vivo*-like setting. Sample sizes, replicates, and statistical methods are specified in the figure legends and the “Statistical analysis” section.

### Human skin and wound specimens

To investigate *in vivo* circRNA expression in human skin wound healing, we collected skin and wound biopsies from 27 healthy volunteers (**Table 1** and **Table S1**). The exclusion criteria for healthy donors were diabetes, skin diseases, unstable heart diseases, infections, bleeding disorders, immune suppression, and any ongoing medical treatments. On the skin of each donor, two or three excisional wounds were created using a 3-mm punch, and the excised skin from these surgical wounds was saved as intact skin control. Wound-edge tissues were collected with a 6-mm punch one day, seven days, and 30 days later. For donors 1-10, full-thickness wound-edge tissues were collected for RNA-seq and qRT-PCR. For donors 11-20, the wound-edge tissues were collected for magnetic cell activation sorting. For donors 21-27, the wound-edge tissues were used for LCM (**Table S1**). Local lidocaine injection was used for anesthesia while sampling. Moreover, skin discarded from plastic surgeries was collected for the establishment of an *ex vivo* wound model (donor 28-30) and the isolation of dermal fibroblasts (donor 31-32) (**Table S1**). Keloid and the surrounding normal skin tissues were collected at the time of surgery from donors 33-40 (**Table S1**).

### RNA-seq library preparation and sequencing

Total RNAs were isolated from the full-depth biopsies of the skin, Wound1, and Wound7 (n = 5/each group), and isolated keratinocytes and fibroblasts from the skin and Wound7 (n = 5/each group) (**Table 1** and **Table S1**) by using the miRNeasy Mini kit (Qiagen, Hilden, Germany) and prepared for library construction. First, the ribosomal RNA (rRNA) was removed using the Epicentre Ribo-zero® rRNA Removal Kit (Epicentre, Road Madison, WI) with a total amount of 2 ug RNA as an input for each library. Second, strand-specific RNA-seq libraries were constructed by using the NEB Next® UltraTM Directional RNA Library Prep Kit for Illumina® (NEB) according to the manufacturer’s instructions. The isolated keratinocytes and fibroblasts RNA-seq libraries were constructed by following the tutorial of the NuGen Ovation Solo RNA-Seq System (Human part no. 0500). Finally, the libraries were sequenced on the Illumina Hiseq 4000 platform (Illumina, Inc., San Diego, CA) by using 150 bp paired-end reads.

### Laser capture microdissection

Frozen tissue samples were cut with a rotary microtome Microm HM355S (ThermoFisher Scientific, Carlsbad, CA) into 10 µm sections and stained with Mayers hematoxylin (HistoLab, Stockholm, Sweden). Laser capture microdissection was performed with Leica LMD7000 (Leica Microsystems, Wetzlar, Germany).

### Magnetic activation cell sorting

Fresh tissue samples were washed 2–3 times in PBS and incubated in 5 U/ml dispase (ThermoFisher Scientific) supplemented with antibiotics (penicillin 50U/I and streptomycin 50 mg/ml. ThermoFisher Scientific) overnight at four °C. The epidermis was separated from the dermis as previously described (*54*). The epidermis was cut into small pieces using scissors and then digested in Trypsin/EDTA Solution (ThermoFisher Scientific) for 15 minutes at 37 °C, from which CD45^-^ cells (mainly composed of keratinocytes) were separated using CD45 Microbeads with MACS MS magnetic columns (Milteney Biotec, North Rhine-Westphalia, Germany). The dermis was incubated in the enzyme mix from the whole skin dissociation kit (Milteney Biotec) for 3 hours according to the manufacturer’s instructions and further processed by Medicon tissue disruptor (BD Biosciences, Stockholm, Sweden). The dermal cell suspension was incubated with CD90 Microbeads, and CD90^+^ fibroblasts were isolated with MACS MS magnetic columns according to the manufacturer’s instructions (Milteney Biotec).

#### *In situ* hybridization

A circGLIS3 probe targeting the circGLIS3 BSJ, a negative control probe targeting Bacillus subtilis dihydrodipicolinate reductase (DapB) gene, and a positive control probe targeting Homo sapiens peptidylprolyl isomerase B (cyclophilin B) (PPIB) mRNA were designed and synthesized by Advanced Cell Diagnostics (ACD, Newark, CA). Human fibroblasts were cultured on slides and fixed in cold 4% formaldehyde for 15 minutes. After dehydration with 50%, 70%, and 100% ethanol, the cells were incubated with Protease III (ACD) at room temperature for 20 min. The slides were then incubated with either a circGLIS3 probe or the negative or positive control probes for two hours at 40°C in HybEZ™ II Hybridization System by using BaseScope™ Reagent Kit v2 – RED Assay (ACD). The hybridization signals were amplified via sequential hybridization of amplifiers and obtained by chromogenic staining with Fast RED dye. Cells were counterstained with 50% hematoxylin for 2 minutes. The cells were visualized with brightfield microscopy on a Nikon eclipse Ni-E microscope (Nikon, Amstelveen, Netherlands) at 20X and 40X magnification.

### Cell culture and functional studies

Human primary dermal fibroblasts, adult (HDFa; Cascade Biologics, Portland, OR) were cultured in Medium 106 (Cascade Biologics) supplemented with 10% Low Serum Growth Supplement (LSGS) and 1% penicillin/ streptomycin at 37°C in 5% CO_2_ (ThermoFisher Scientific).

Dermal fibroblasts were isolated from adult human skin from abdominal or thigh reduction plastic surgery (n = 2) (donors 31 and 32 in **Table S1**). Six-mm full-depth skin biopsies were collected with a punch knife and washed with PBS. The tissues were then placed in a culture plate and left to attach to the bottom. Dulbecco’s Modified Eagle Medium (DMEM) supplemented with 10% heat-inactivated fetal bovine serum (FBS) and 1% penicillin-streptomycin (ThermoFisher Scientific) was then added to the culture plate, which was kept at 37 °C in 5% CO2. Fibroblasts grew out from the tissues, and the culture became confluent in approximately two weeks. Cells were passaged once for expansion. Passage two fibroblasts were cryopreserved. Fibroblasts in passages three and four were used in this study.

NIH/3T3 mouse embryonic fibroblasts (CRL-1658™; ATCC, Manassas, VA) were cultured in DMEM medium supplemented with 10% heat-inactivated FBS and 1% penicillin-streptomycin at 37°C in 5% CO_2_ (ThermoFisher Scientific). HEK293T cells were cultured in DMEM medium supplemented with 10% FCS and 1% penicillin-streptomycin at 37°C in 5% CO_2_.

To evaluate RNA stability, we incubated human fibroblasts with Actinomycin-D 5 µg/ml for up to 24 hours. To study the mechanism regulating circGLIS3 expression, we treated human fibroblasts with IL-1α (20 ng/ml), IL-6 (50 ng/ml), IL-8 (50 ng/ml), IL-22 (30 ng/ml), IL-36α (100 ng/ml), TNF-α (50 ng/ml), TGF-β1 (20 ng/ml), TGF-β2 (10 ng/ml), TGF-β3 (20 ng/ml), BMP-2 (100 ng/ml), EGF (20 ng/ml), IGF-1 (20 ng/ml), FGF-2 (30 ng/ml), VEGFA (20 ng/ml), HB-EGF (20 ng/ml) or PBS as control for 24 hours and circGLIS3 expression was analyzed by qRT-PCR. All these cytokines and growth factors were purchased from either ImmunoTools (Frieshoyte, Germany) or R&D Systems (Minneapolis, MN) (**Table S3**).

To study the functions of circGLIS3 in fibroblasts, we knocked down or overexpressed circGLIS3. For knockdown experiments, cells at 60–70% confluence were transfected with 10 nM of siRNA targeting circGLIS3 or a scrambled siRNA for 24 and 48 hours using Lipofectamine™ 3000 (ThermoFisher Scientific). To overexpress circGLIS3, we transfected fibroblasts at 80-90% confluence with circGLIS3 overexpression plasmid (pLC5-circGLIS3) or mock vector (pLC5-empty) (250ng/ml) with Lipofectamine™ 3000 for 48 hours. The successful modulation of circGLIS3 expression level was confirmed by qRT-PCR (**Fig. 3E, F**, and **Fig. S2**). For evaluation of the effect of circGLIS3 on TGF-β signaling, human fibroblasts with either circGLIS3 depletion or overexpression or 3T3 cells with circGlis3 knockdown that had been transfected for 24h were stimulated with 5 ng/ml TGF-β1 (R&D Systems) for 24 hours. The corresponding amount of TGF-β1 reconstitution buffer was used as vehicle negative control. These cells were used for qRT-PCR analysis. To test the endogenous PCPE-1 protein turnover, we treated human fibroblasts with 0.5 µM MG132 (Sigma-Aldrich, Cat. No. M7449) and 5 µg/ml cycloheximide (Sigma-Aldrich, Cat No. C4859).

### Luciferase assay

To evaluate the effect of circGLIS3 on the responsiveness of human fibroblasts to TGF-β1 stimulation, we used a TGF-β reporter plasmid pSBE4-Luc (Addgene, plasmid #16495). This plasmid contains four tandem copies of the Smad binding sites, which drive the transcription of the Firefly luciferase reporter gene (*27*). pBV-Luc (Addgene, plasmid #16539), a luciferase reporter plasmid with very low basal activity, was used as a negative control (*55*). Human dermal fibroblasts were co-transfected with the luciferase reporters (200 ng/ml), together with 10 nM siRNA targeting circGLIS3 or scrambled control, using the Lipofectamine™ 3000 (ThermoFisher Scientific). One day later, the transfected cells were treated with 5 ng/ml TGF-β1 (R&D Systems) for 24 hours. Luciferase activity was analyzed using the Dual-Luciferase® Reporter Assay System and read with GloMax®-Multi Detection System (Promega, Madison, WI).

### Plasmids construction

The circGLIS3 overexpression plasmid was constructed with the help of Guangzhou Geneseed Biotech Co. (Guangzhou, China). In brief, the pLC5-ciR vector, which includes front and back circular frames for the circularization of the transcripts, was used as the backbone plasmid. The front circular frame contains an endogenous flanking genomic sequence with the EcoRI restriction site, and the back circular frame contains part of the inverted upstream sequence with the BamHI restriction site. The cDNA encoding circGLIS3 in HEK293T cells was amplified using the primers listed in **Table S3**. The amplicon, which contained an EcoRI site, the circGLIS3 linear sequence with the corresponding splice sites, and a BamHI site, was then cloned into the pLC5-ciR backbone vector between the two frames. Vector construction was verified by Sanger sequencing. A mock vector containing only a nonsense sequence between the two circular frames was used as a control plasmid.

The circGLIS3 overexpression plasmids with or without MS2 hairpins were constructed with the help of Creative Biogene Biotechnology (Shirley, NY). In brief, the cDNA encoding circGLIS3 or circGLIS3-MS2 were subcloned into the pLO-circRNA backbone by restriction digestion with EcoRI and BamHI and ligation with T4 DNA ligase. Vector construction was verified by Sanger sequencing, and the primers used are listed in **Table S3**.

### MS2-mediated pulldown of circGLIS3-bound proteins and mass spectrometry

Pulldown of MS2-tagged circGLIS3 and its protein interactome was performed using previously published methods (*56, 57*). In brief, we co-transfected HEK293T cells with 1 µg circGLIS3 overexpression plasmids with or without the MS2 hairpins (circGLIS3 and circGLIS3-MS2) together with a captured protein expression plasmid (MS2-CP) containing a FLAG tag for 48 hours by using Lipofectamine™ Reagent and PLUS™ reagent (ThermoFisher Scientific) (**Fig. 4A**). The immunoprecipitation of the circGLIS3-RBP complex was performed with protein A+G beads coated with an anti-FLAG antibody. RNA-protein complexes were eluted from the beads; RNA and protein fractions were isolated. The enrichment of circGLIS3 in the circGLIS3-MS2 group after immunoprecipitation was validated by qRT-PCR. The protein fractions from circGLIS3-MS2 (test) and circGLIS3 (control) were analyzed with mass spectrometry. Briefly, the proteins in the eluate were reduced with 0.05M TCEP solution at 60°C for 1 hour and then alkylated with 55 nM MMTS for 45 min at room temperature. The solution was filtered on 10 kDa centrifugal filter devices for 20 min at 12000 x g. The proteins were then digested with trypsin at 37°C overnight using an enzyme-to-protein ratio of 1:50. The resulting peptides were collected by centrifugation and vacuum dried at low temperature. Peptides were then dissolved in 2% ACN and 0.1% formic acid and analyzed on a Thermo Scientific Q Exactive Mass Spectrometer (ThermoFisher Scientific). The MS was operated in data-dependent mode, automatically switching between MS and MS2 acquisition, with a mass resolution of 70,000 and 17,500, respectively. Mascot software was used for protein identification. MS raw files were searched against a database of 20386 *Homo sapiens* sequences from uniprot.org. Protein scores (**Table S4**) were derived from ion scores where individual ion scores > 22 indicated identity or extensive homology (p < 0.05). The protein lists identified in each group were overlapped with a list of 924 proteins with enriched expression in human skin fibroblasts (**Table S5**) obtained from proteinatlas.org (**Fig. 4C**).

### RNA-binding protein immunoprecipitation

To test the interaction between human circGLIS3 and PCPE-1, we performed a RIP assay using Magna RIP RNA-Binding Protein Immunoprecipitation Kit (Millipore, Burlington, MA). HDFa were lysed in RIP lysis buffer, and then 100 μl of whole cell extract was incubated with anti-human PCPE-1 antibody (sc-73002, Santa Cruz Biotechnology) coated on A + G magnetic beads (Millipore) in RIP buffer. Normal rabbit IgG (Millipore) was used as a negative control. The samples were then incubated with proteinase K to digest protein, and the immunoprecipitated RNA was isolated. circGLIS3 and *GAPDH* mRNA levels were detected by qRT-PCR.

### Cellular thermal shift assay (CETSA) and immunoblot

Human dermal fibroblasts transfected with either 10 nM si-circGLIS3 or scrambled siRNA (Eurofins Genomics, Ebersberg, Germany) for 48 hours and treated with 5 ng/ml TGF-β1 (R&D Systems) were rinsed and pelleted in PBS. Cells were resuspended in PBS supplemented with protease inhibitors (Roche, Basel, Switzerland) and counted to normalize equal cell density between conditions. Cells were lysed with three rounds of freeze-thaw cycles by incubating for three minutes on dry ice and three minutes in a water bath at 37°C, followed by centrifugation at 16,000 × g for 20 minutes at 4°C. The supernatant was then aliquoted into PCR tubes and heated individually at different temperatures (range: 55-90°C, 2.5°C increments) for 3 minutes in a gradient thermal cycler ProFlex PCR System (ThermoFisher Scientific) and immediately cooled down at room temperature. After centrifugation (20,000 × g for 20 minutes at 4°C), the supernatant was transferred to a new tube and prepared for immunodetection with an anti-human PCPE-1 antibody (1:25; catalog sc-73002; Santa Cruz Biotechnology) with Protein Simple Jess/Wes capillary-based system (Bio-Techne, Minneapolis, MN) according to manufacturer instructions.

### Human *ex vivo* wound model

To evaluate the effect of circGLIS3 in a physiologically relevant model of human skin wound healing, we employed a human *ex vivo* wound model (*58*). Human skin was obtained from abdominal reduction surgeries or thighplasty (donors 28-30 in **Table S1**). The wounds were made using a 2 mm biopsy punch on the epidermal side of the skin (2-4 wounds per donor), excised using a 6 mm biopsy punch, and subsequently transferred to a cell culture plate containing DMEM plus 10% FBS and antibiotics (penicillin 50 U/l and streptomycin 50 mg/ml; ThermoFisher Scientific). MaxSuppressor In Vivo RNA-LANCEr II (Bioo Scientific, Austin, TX) was mixed with 0.1 nmol siRNA targeting circGLIS3 or a scrambled siRNA (Eurofins Genomics) in a volume of 5 µl per wound. The siRNA-lipid complexes were mixed 1:2 (volumes) in 30% pluronic F-127 gel (Sigma-Aldrich). 15 µl mixture was topically applied on the wounds immediately after injury and 3 days later. Wound samples were collected one day later for gene expression analysis (**Fig. 6D, E**) and six days after injury for histological analysis (**Fig. 6B, C, F, G**).

### Statistics

Data analysis was performed using R and GraphPad 8.4.0 (GraphPad Software). All quantitative data were presented as means ± SD. Normality and distribution of data were checked with the Shapiro-Wilk test (p < 0.05 indicated data that did not pass the normality test). Comparison between two groups was performed with a two-tailed Student’s t-test (parametric) or Mann-Whitney U test (non-parametric, unpaired), or Wilcoxon test (non-parametric, paired). Comparison between more than two groups that contained paired data (matched samples or repeated measures) was made with RM one-way ANOVA and Tukey’s multiple comparisons test (parametric data) or Friedman test and Dunn’s multiple comparisons test (non-parametric data). Comparison between more than two groups with unpaired data was performed with Ordinary one-way ANOVA and Dunnett’s multiple comparisons test (parametric data) or Kruskal-Wallis and Dunn’s multiple comparisons test (non-parametric data). p-value < 0.05 was considered statistically significant.

## Supplementary Materials

### Materials and Methods

Fig. S1. The molecular characteristics of circGLIS3.

Fig. S2. Modulation of circGLIS3 levels in human dermal fibroblasts.

Fig. S3. circGlis3 regulates Tgf-β1 target genes in mouse fibroblasts.

Table S1. Human sample information.

Table S2. Differentially expressed circRNAs in human day-7 wounds compared to the matched skin (tissue biopsies, dermal fibroblasts, and epidermal keratinocytes).

Table S3. List of reagents used in this study.

Table S4. Mass spectrometry analysis of protein interactome of circGLIS3.

Table S5. Proteins expressed in human skin fibroblasts.

## Acknowledgments

We thank all the tissue donors participating in this study and Helena Griehsel for technical support during sample collection. We thank the Microarray core facility at Novum, BEA, which is supported by the board of research at KI and the research committee at the Karolinska hospital. The computations/data handling was enabled by resources in projects of sens2020010 and SNIC2019/8-262 provided by the Swedish National Infrastructure for Computing (SNIC) at UPPMAX, partially funded by the Swedish Research Council through grant agreement no. 2018-05973.

## Funding

Swedish Research Council (Vetenskapsrådet) grants 2016-02051 and 2020-01400 (NXL)

Ragnar Söderbergs Foundation, grant M31/15 (NXL)

Welander and Finsens Foundation (Hudfonden) (NXL)

Ming Wai Lau Centre for Reparative Medicine (NXL)

LEO Foundation (NXL)

Cancerfonden (NXL)

Karolinska Institutet (NXL)

## Author contributions

Conceptualization: NXL, MAT, PS Methodology: MAT, ZL, MV, MP, AW, DL, PS

Investigation: MAT, QW, MV, MP, ZL, LZ, DL, GN, JG, YX, XB, PS

Visualization: MAT, NXL, GN Funding acquisition: NXL

Project administration: MAT, NXL Supervision: NXL, PS

Writing – original draft: MAT, NXL

Writing – review & editing: MAT, QW, DL, YX, GN, JG, MV, MP, ZL, LZ, XB, AW, PS, NXL

## Data and materials availability

The circRNA expression data from RNA-seq of human skin and wound tissues have been previously reported (*13*) and presented as a web resource at https://www.xulandenlab.com/humanwounds-circrna. The circRNA expression data from RNA-seq of isolated keratinocytes and fibroblasts are presented in the supplementary material in Table S2. The transcriptomic profiling of dermal fibroblasts with circGLIS3 knockdown by microarray has been deposited to Gene Expression Omnibus with the ascension number GSE196260 (token: adkficmuhnmdfsv). All codes required to reanalyze the data reported in this paper can be requested from the lead contact.

## REFERENCES AND NOTES

1. K. Jarbrink et al., The humanistic and economic burden of chronic wounds: a protocol for a systematic review. Syst Rev 6, 15 (2017).

2. R. G. Frykberg, J. Banks, Challenges in the Treatment of Chronic Wounds. Adv Wound Care (New Rochelle) 4, 560–582 (2015).

3. G. G. Gauglitz, H. C. Korting, T. Pavicic, T. Ruzicka, M. G. Jeschke, Hypertrophic scarring and keloids: pathomechanisms and current and emerging treatment strategies. Mol Med 17, 113–125 (2011).

4. J. M. Reinke, H. Sorg, Wound repair and regeneration. Eur Surg Res 49, 35–43 (2012).

5. K. S. Midwood, L. V. Williams, J. E. Schwarzbauer, Tissue repair and the dynamics of the extracellular matrix. Int J Biochem Cell Biol 36, 1031–1037 (2004).

6. W. R. Jeck et al., Circular RNAs are abundant, conserved, and associated with ALU repeats. RNA 19, 141–157 (2013).

7. S. Memczak et al., Circular RNAs are a large class of animal RNAs with regulatory potency. Nature 495, 333–338 (2013).

8. X. Li, L. Yang, L. L. Chen, The Biogenesis, Functions, and Challenges of Circular RNAs. Mol Cell 71, 428–442 (2018).

9. L. L. Chen, The expanding regulatory mechanisms and cellular functions of circular RNAs. Nat Rev Mol Cell Biol 21, 475–490 (2020).

10. Q. Yang, F. Li, A. T. He, B. B. Yang, Circular RNAs: Expression, localization, and therapeutic potentials. Mol Ther 29, 1683–1702 (2021).

11. X. Mao, Y. Cao, Z. Guo, L. Wang, C. Xiang, Biological roles and therapeutic potential of circular RNAs in osteoarthritis. Mol Ther Nucleic Acids 24, 856–867 (2021).

12. L. Santer, C. Bar, T. Thum, Circular RNAs: A Novel Class of Functional RNA Molecules with a Therapeutic Perspective. Mol Ther 27, 1350–1363 (2019).

13. M. A. Toma et al., Circular Rna Signatures Of Human Healing And Non-Healing Wounds. J Invest Dermatol, (2022).

14. T. Zhang et al., Current potential therapeutic strategies targeting the TGF-beta/Smad signaling pathway to attenuate keloid and hypertrophic scar formation. Biomed Pharmacother 129, 110287 (2020).

15. H. P. Ehrlich et al., Morphological and immunochemical differences between keloid and hypertrophic scar. Am J Pathol 145, 105–113 (1994).

16. W. Wu, P. Ji, F. Zhao, CircAtlas: an integrated resource of one million highly accurate circular RNAs from 1070 vertebrate transcriptomes. Genome Biol 21, 101 (2020).

17. S. Dodbele, N. Mutlu, J. E. Wilusz, Best practices to ensure robust investigation of circular RNAs: pitfalls and tips. EMBO Rep 22, e52072 (2021).

18. T. Gutschner, M. Hammerle, S. Diederichs, MALAT1 -- a paradigm for long noncoding RNA function in cancer. J Mol Med (Berl) 91, 791–801 (2013).

19. S. Bandiera et al., Nuclear outsourcing of RNA interference components to human mitochondria. PLoS One 6, e20746 (2011).

20. O. Bensaude, Inhibiting eukaryotic transcription: Which compound to choose? How to evaluate its activity? Transcription 2, 103–108 (2011).

21. H. Suzuki, T. Tsukahara, A view of pre-mRNA splicing from RNase R resistant RNAs. Int J Mol Sci 15, 9331–9342 (2014).

22. W. R. Jeck, N. E. Sharpless, Detecting and characterizing circular RNAs. Nat Biotechnol 32, 453–461 (2014).

23. S. Barrientos, O. Stojadinovic, M. S. Golinko, H. Brem, M. Tomic-Canic, Growth factors and cytokines in wound healing. Wound Repair Regen 16, 585–601 (2008).

24. J. W. Penn, A. O. Grobbelaar, K. J. Rolfe, The role of the TGF-beta family in wound healing, burns and scarring: a review. Int J Burns Trauma 2, 18–28 (2012).

25. M. B. Vaughan, E. W. Howard, J. J. Tomasek, Transforming growth factor-beta1 promotes the morphological and functional differentiation of the myofibroblast. Exp Cell Res 257, 180–189 (2000).

26. S. Bhattacharyya et al., Tenascin-C drives persistence of organ fibrosis. Nat Commun 7, 11703 (2016).

27. L. Zawel et al., Human Smad3 and Smad4 are sequence-specific transcription activators. Mol Cell 1, 611–617 (1998).

28. Q. Zhu et al., Synergistic effect of PCPE1 and sFRP2 on the processing of procollagens via BMP1. FEBS Lett 593, 760 (2019).

29. E. V. Rusilowicz-Jones, S. Urbe, M. J. Clague, Protein degradation on the global scale. Mol Cell 82, 1414–1423 (2022).

30. R. Jafari et al., The cellular thermal shift assay for evaluating drug target interactions in cells. Nat Protoc 9, 2100–2122 (2014).

31. L. E. Tracy, R. A. Minasian, E. J. Caterson, Extracellular Matrix and Dermal Fibroblast Function in the Healing Wound. Adv Wound Care (New Rochelle) 5, 119–136 (2016).

32. M. Pakyari, A. Farrokhi, M. K. Maharlooei, A. Ghahary, Critical Role of Transforming Growth Factor Beta in Different Phases of Wound Healing. Adv Wound Care (New Rochelle) 2, 215–224 (2013).

33. J. Lu et al., Increased expression of latent TGF-beta-binding protein 4 affects the fibrotic process in scleroderma by TGF-beta/SMAD signaling. Lab Invest 97, 591–601 (2017).

34. M. K. Lichtman, M. Otero-Vinas, V. Falanga, Transforming growth factor beta (TGF-beta) isoforms in wound healing and fibrosis. Wound Repair Regen 24, 215–222 (2016).

35. X. Li, Z. He, J. Zhang, Y. Han, Identification of crucial non-coding RNAs and mRNAs in hypertrophic scars via RNA sequencing. FEBS Open Bio 11, 1673–1684 (2021).

36. Y. Li et al., CircGLIS3 Promotes High-Grade Glioma Invasion via Modulating Ezrin Phosphorylation. Front Cell Dev Biol 9, 663207 (2021).

37. S. Wu et al., Circular RNA circGLIS3 promotes bladder cancer proliferation via the miR-1273f/SKP1/Cyclin D1 axis. Cell Biol Toxicol 38, 129–146 (2022).

38. Z. Wu, H. Jiang, H. Fu, Y. Zhang, A circGLIS3/miR-644a/PTBP1 positive feedback loop promotes the malignant biological progressions of non-small cell lung cancer. Am J Cancer Res 11, 108–122 (2021).

39. J. Xu et al., Overexpression of hsa_circ_0002874 promotes resistance of non-small cell lung cancer to paclitaxel by modulating miR-1273f/MDM2/p53 pathway. Aging (Albany NY) 13, 5986–6009 (2021).

40. L. Xiong et al., Lipotoxicity-induced circGlis3 impairs beta cell function and is transmitted by exosomes to promote islet endothelial cell dysfunction. Diabetologia 65, 188–205 (2022).

41. R. Adar, E. Kessler, B. Goldberg, Evidence for a protein that enhances the activity of type I procollagen C-proteinase. Coll Relat Res 6, 267–277 (1986).

42. E. Hassoun, M. Safrin, H. Ziv, S. Pri-Chen, E. Kessler, Procollagen C-Proteinase Enhancer 1 (PCPE-1) as a Plasma Marker of Muscle and Liver Fibrosis in Mice. PLoS One 11, e0159606 (2016).

43. G. Kessler-Icekson, H. Schlesinger, S. Freimann, E. Kessler, Expression of procollagen C-proteinase enhancer-1 in the remodeling rat heart is stimulated by aldosterone. Int J Biochem Cell Biol 38, 358–365 (2006).

44. V. W. Wong, F. You, M. Januszyk, G. C. Gurtner, A. A. Kuang, Transcriptional profiling of rapamycin-treated fibroblasts from hypertrophic and keloid scars. Ann Plast Surg 72, 711–719 (2014).

45. L. Ma et al., Comparative proteomic analysis of extracellular matrix proteins secreted by hypertrophic scar with normal skin fibroblasts. Burns Trauma 2, 76–83 (2014).

46. A. E. Postlethwaite et al., Modulation of fibroblast functions by interleukin 1: increased steady-state accumulation of type I procollagen messenger RNAs and stimulation of other functions but not chemotaxis by human recombinant interleukin 1 alpha and beta. The Journal of cell biology 106, 311–318 (1988).

47. A. Tang, B. A. Gilchrest, Regulation of keratinocyte growth factor gene expression in human skin fibroblasts. J Dermatol Sci 11, 41–50 (1996).

48. Y. Kawaguchi, IL-1 alpha gene expression and protein production by fibroblasts from patients with systemic sclerosis. Clin Exp Immunol 97, 445–450 (1994).

49. T. Z. Kirk, M. D. Mayes, IL-1 rescues scleroderma myofibroblasts from serum-starvation-induced cell death. Biochem Biophys Res Commun 255, 129–132 (1999).

50. A. Wang et al., Circular RNA hsa_circ_0084443 Is Upregulated in Diabetic Foot Ulcer and Modulates Keratinocyte Migration and Proliferation. Adv Wound Care (New Rochelle) 9, 145–160 (2020).

51. D. Han, W. Liu, G. Li, L. Liu, Circ_PRKDC knockdown promotes skin wound healing by enhancing keratinocyte migration via miR-31/FBN1 axis. J Mol Histol 52, 681–691 (2021).

52. L. N. Jiang, X. Ji, W. Liu, C. Qi, X. Zhai, Identification of the circ_PRKDC/miR-20a-3p/RASA1 axis in regulating HaCaT keratinocyte migration. Wound Repair Regen, (2021).

53. Z. G. Yang et al., The Circular RNA Interacts with STAT3, Increasing Its Nuclear Translocation and Wound Repair by Modulating Dnmt3a and miR-17 Function. Mol Ther 25, 2062–2074 (2017).

54. S. Cheuk et al., CD49a Expression Defines Tissue-Resident CD8(+) T Cells Poised for Cytotoxic Function in Human Skin. Immunity 46, 287–300 (2017).

55. T. C. He, T. A. Chan, B. Vogelstein, K. W. Kinzler, PPARdelta is an APC-regulated target of nonsteroidal anti-inflammatory drugs. Cell 99, 335–345 (1999).

56. J. H. Yoon, M. Gorospe, Identification of mRNA-Interacting Factors by MS2-TRAP (MS2-Tagged RNA Affinity Purification). Methods Mol Biol 1421, 15–22 (2016).

57. L. M. Holdt et al., Circular non-coding RNA ANRIL modulates ribosomal RNA maturation and atherosclerosis in humans. Nat Commun 7, 12429 (2016).

58. O. Stojadinovic, M. Tomic-Canic, Human ex vivo wound healing model. Methods Mol Biol 1037, 255–264 (2013).

