## Supplementary Material - Material and Methods, Supplementary Figure 1, 2 and 3 and Supplementary Table 1 and 3 for "The injury-induced circular RNA circGLIS3 activates dermal fibroblasts to promote wound healing"

**The PDF file includes:**

Materials and Methods

Figures. S1 to S3

Tables S1 and S3

**Other Supplementary Material for this manuscript includes the following:**

Tables S2, S4, S5

### MATERIALS AND METHODS

#### CircRNA identification and differential expression analysis of RNA-seq data

CircRNA identification and differential expression (DE) analysis of RNA-seq data was performed as previously reported (*1*) and presented as a browsable web resource (https://www.xulandenlab.com/humanwounds-circrna). Briefly, raw reads were first filtered using the Trimmomatic v0.36 package (*2*) to remove adaptor sequences and low-quality bases. Clean reads were then mapped to the human reference genome (GRCh38.p12) with the GENCODE genes annotation (version 31) using STAR v2.7.1 (*3*). The unmapped chimeric reads were used for circRNA identification using DCC software (*4*). CircRNAs were filtered by at least two back-spliced junction reads in a minimum of two samples. The expression of circRNAs was normalized to FPM (fragments mapped to back-splicing junctions per million mapped fragments) as previously described (*5*). CircRNAs were annotated based on chromosomal location and the overlap with three circRNA databases: circAtlas v2 (*6*), circBase (*7*), and CIRCpedia v2 (*8*). Differential circRNA analysis was performed by using the DESeq2 package (*9*). P-values calculated from the Wald test were adjusted by performing Benjamini-Hochberg (BH) multiple testing to estimate the false discovery rate (FDR). The differentially expressed circRNAs were defined as fold change greater than 2 and P value $< 0.05$. The most abundant DE circRNAs were selected based on mean normalized read count values higher than 1 (**Fig. 1B, Table S2**).

#### Mouse skin wound specimens

To investigate the expression of circGlis3 in mouse wounds, we collected skin and wound biopsies from 15 C57BL/6 mice. All the mice were housed individually for one week before the surgery and during the experiment. General anesthesia was performed with 3% isoflurane

(Abbott, Chicago, IL). During the first 2 days, the animals received subcutaneous buprenorphine (30 μg/kg) twice a day to relieve any possible distress caused by the experimental procedure. The hair of the back was shaved with a shaver followed by a depilatory cream. On the same day, two 4-mm full-thickness wounds were made on the back using a biopsy punch. The excised skin was saved. The wound-edges were excised with a 6 mm-biopsy punch at day 3 (n = 5; two wounds/mouse), day 7 (n = 5; one wound/mouse), and day 10 (n = 5; one wound/mouse) after wounding. The animals were sacrificed before tissue collection. Biopsies were used for RNA extraction and qRT-PCR. The animal experiment procedure was reviewed and approved by the North Stockholm Ethical Committee for Care and Use of Laboratory Animals (Stockholm, Sweden).

**RNA extraction and qRT-PCR**

Full-thickness skin and wound biopsies were homogenized using TissueLyser LT (Qiagen, Hilden, Germany) before RNA extraction. Total RNA was extracted from human tissue and microdissected tissue and isolated cells using the miRNeasy Mini kit (Qiagen) and from cultured cells using TRIzol reagent (ThermoFisher Scientific). Gene expression was either determined by human or mouse TaqMan expression assays (ThermoFisher Scientific) (ACTA2, COL1A1, COL4A1, and FN1) or by SybrGreen expression assays (ThermoFisher Scientific) (GLIS3 and circGLIS3) and normalized based on the values of the housekeeping gene 18S, B2M, ACTB, or GAPDH. The information for all the primers/probes used in this study is listed in **Table S3**.

**Cell fractionation**

Cytoplasm and nucleus of dermal fibroblasts were separated by using Nuclear Extract Kit (Active Motif) following the manufacturer's instructions. Mitochondria were isolated using Mitochondria Isolation Kit for Cultured Cells (ThermoFisher Scientific). RNA was extracted from these fractions using Trizol (ThermoFisher Scientific). qRT-PCR was performed to analyze the expression of circGLIS3, GAPDH, MALAT1, and 16S rRNA.

**PCR and Sanger sequencing**

To confirm the circularity of both human and mouse circGLIS3, we performed PCR with outward-facing primers to detect head-to-tail splice sites of circGLIS3. Moreover, fibroblast RNA was digested with 1 unit RNaseR (Lucigen, Biosearch Technologies, Middleton, WI) per 1 μg RNA to enrich for circRNAs. Fifty ng cDNA obtained from human or mouse fibroblasts (3T3 cells) were amplified with the divergent primers (**Table S3**) by using PCR Master Mix (2X) (ThermoFisher Scientific) for 33 cycles. The PCR products were then analyzed by DNA electrophoresis using E-Gel™ Agarose Gels with SYBR™ Safe DNA Gel Stain, 2% (Invitrogen, Waltham, MA, USA). The bands with expected sizes were cut out, and the DNA was purified using QIAquick Gel Extraction Kit (Qiagen). Purified DNA was then analyzed by Sanger sequencing using ABI 3730 PRISM® DNA Analyzer at the KIgene core facility at Karolinska Institutet (Stockholm, Sweden).

**Gene expression microarray and analysis**

Expression profiling of human fibroblasts transfected with either scrambled siRNA or siRNA targeting circGLIS3 for 24 hours (n = 3 per group) was carried out by using the human Clariom™ S assay (ThermoFisher Scientific) at the Bioinformatics and Expression Analysis (BEA) core facility at Karolinska Institutet. In brief, total RNA was extracted using TRIzol, and RNA quality and quantity were determined using Agilent 2200 Tapestation with RNA ScreenTape and Nanodrop 1000. 150 nanograms of total RNA were used to prepare cDNA following the GeneChip WT PLUS Reagent Kit labeling protocol. Standardized array processing procedures recommended by Affymetrix, including hybridization, fluidics processing, and scanning, were used.

Expression data were analyzed by using Transcriptome Analysis Console 4.0. Genes showing at least 1.5-fold regulation and FDR less than 0.05 were considered as differentially expressed. MetaCore software (Thomson Reuters, Toronto, Canada) was used for gene ontology analyses to identify the top enriched biological pathways and processes among the genes regulated by circGLIS3. Gene set enrichment analysis (GSEA) was performed to evaluate the enrichment of TGF-β1 target genes (obtained from GSE79621 (*10*), **Table S3**) among the genes regulated by circGLIS3 in fibroblasts by using public software from Broad Institute. The data reported here have been deposited to Gene Expression Omnibus with the ascension number GSE196260 (with a secure token 'adkficmuhnmdfsv' to allow the review of the data).

**Western blot**

Fibroblasts protein lysates were extracted by using radioimmunoprecipitation assay (RIPA) buffer (ThermoFisher Scientific) and analyzed by Western blotting with the following antibodies: anti-human COL1A (1:1000; catalog sc-59772; Santa Cruz Biotechnology, Dallas, TX), anti-human PCPE-1 (1:1000; catalog sc-73002; Santa Cruz Biotechnology), and anti-human α-SMA (1:1000; catalog MAB1420; R&D Systems). B-actin expression was visualized using an HRP-coupled anti-human actin antibody (1:20,000; catalog A3854; Sigma-Aldrich, St. Louis, MO).

**Immunofluorescent staining**

Formalin-fixed paraffin-embedded tissue samples of human *ex vivo* wounds were deparaffinized and subjected to heat-induced antigen retrieval in Tris-EDTA buffer (pH = 9) at 98°C for 25 minutes. The tissue sections were blocked with 5% bovine serum albumin (BSA; ThermoFisher Scientific) and stained with α-smooth muscle actin (1:100; MAB1420, R&D systems) at 4°C overnight. Sections were then washed with TBST (TBS with 0.1% Triton-X) and incubated with highly pre-cross absorbed Alexa Fluor 555 donkey anti-mouse IgG secondary antibody (1:200 in TBST; A31570, Invitrogen, Waltham, MA). After being washed with TBST, sections were mounted with ProLong™ Diamond Antifade Mountant with DAPI (ThermoFisher Scientific).

Human dermal fibroblasts were fixed with 4% paraformaldehyde in PBS, blocked with 2.5% BSA in PBS, followed by permeabilization with 0.1% Triton-X in PBS. Cells were stained with primary antibodies against α-smooth muscle actin (MAB1420, R&D systems) or procollagen C-proteinase enhancer-1 (PCPE-1; sc-73002, Santa Cruz Biotechnology) in 1:100 dilution in a blocking solution. Highly pre-cross absorbed Alexa Fluor 555 donkey anti-mouse IgG was used as a secondary antibody (1:200; A31570, Invitrogen), and cells were mounted with ProLong™ Diamond Antifade Mountant with DAPI (ThermoFisher Scientific). Immunofluorescence-stained cells and tissue sections were examined using a Nikon eclipse Ni-E fluorescence microscope (Nikon) at 20X magnification.

### SUPPLEMENTAL REFERENCES


1. M. A. Toma *et al.*, Circular Rna Signatures Of Human Healing And Non-Healing Wounds. *J Invest Dermatol*, (2022).

2. A. M. Bolger, M. Lohse, B. Usadel, Trimmomatic: a flexible trimmer for Illumina sequence data. *Bioinformatics* **30**, 2114-2120 (2014).
3. A. Dobin *et al.*, STAR: ultrafast universal RNA-seq aligner. *Bioinformatics* **29**, 15-21 (2013).
4. Y. Liao, G. K. Smyth, W. Shi, featureCounts: an efficient general purpose program for assigning sequence reads to genomic features. *Bioinformatics* **30**, 923-930 (2014).
5. J. Cheng, F. Metge, C. Dieterich, Specific identification and quantification of circular RNAs from sequencing data. *Bioinformatics* **32**, 1094-1096 (2016).
6. W. Wu, P. Ji, F. Zhao, CircAtlas: an integrated resource of one million highly accurate circular RNAs from 1070 vertebrate transcriptomes. *Genome Biol* **21**, 101 (2020).
7. P. Glazar, P. Papavasileiou, N. Rajewsky, circBase: a database for circular RNAs. *RNA* **20**, 1666-1670 (2014).
8. R. Dong, X. K. Ma, G. W. Li, L. Yang, CIRCpedia v2: An Updated Database for Comprehensive Circular RNA Annotation and Expression Comparison. *Genomics Proteomics Bioinformatics* **16**, 226-233 (2018).
9. M. I. Love, W. Huber, S. Anders, Moderated estimation of fold change and dispersion for RNA-seq data with DESeq2. *Genome Biol* **15**, 550 (2014).
10. S. Bhattacharyya *et al.*, Tenascin-C drives persistence of organ fibrosis. *Nat Commun* **7**, 11703 (2016).

### SUPPLEMENTAL FIGURES

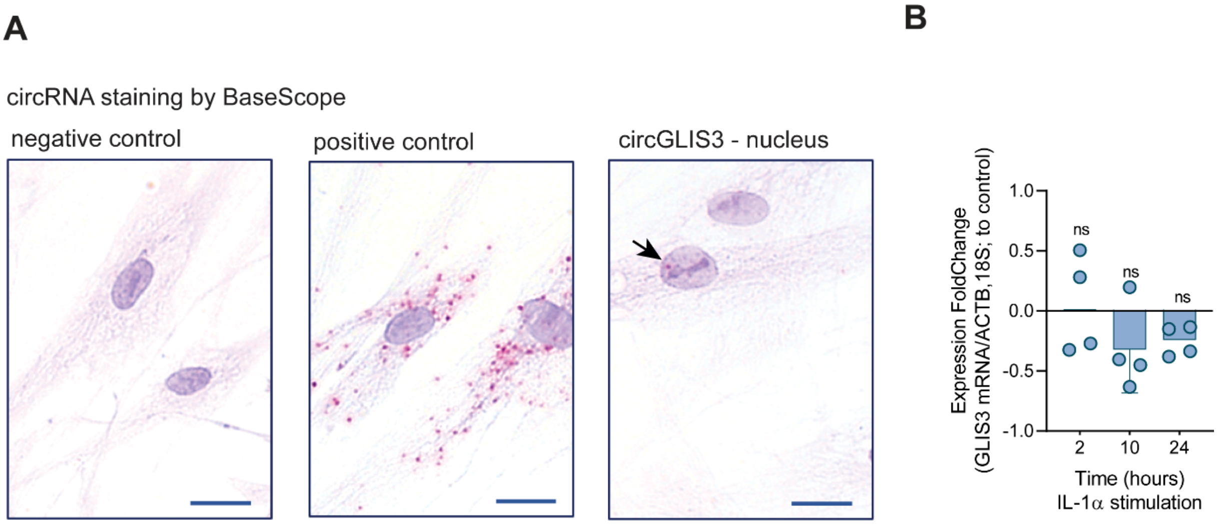


**Figure S1. The molecular characteristics of circGLIS3.** (**A**) RNA *in situ* hybridization using BaseScope™ assay in human dermal fibroblasts. The pictures show the signal from the Fast RED dye of the negative control probe (detecting DapB), the positive control probe (detecting *PPIB* mRNA), and the probe targeting circGLIS3 BSJ. The black arrow indicates the circGLIS3 signal in the nucleus. (**B**) qRT-PCR analysis of *GLIS3* mRNA expression in fibroblasts treated with IL-1α (20 ng/μl) for 2-24 hours (n=4). Data show *GLIS3* mRNA expression normalized to housekeeping genes *ACTB* and *18S* and relative to the levels in the untreated cells. ns = not significant, P value > 0.05 by two-tailed Student's t-test.

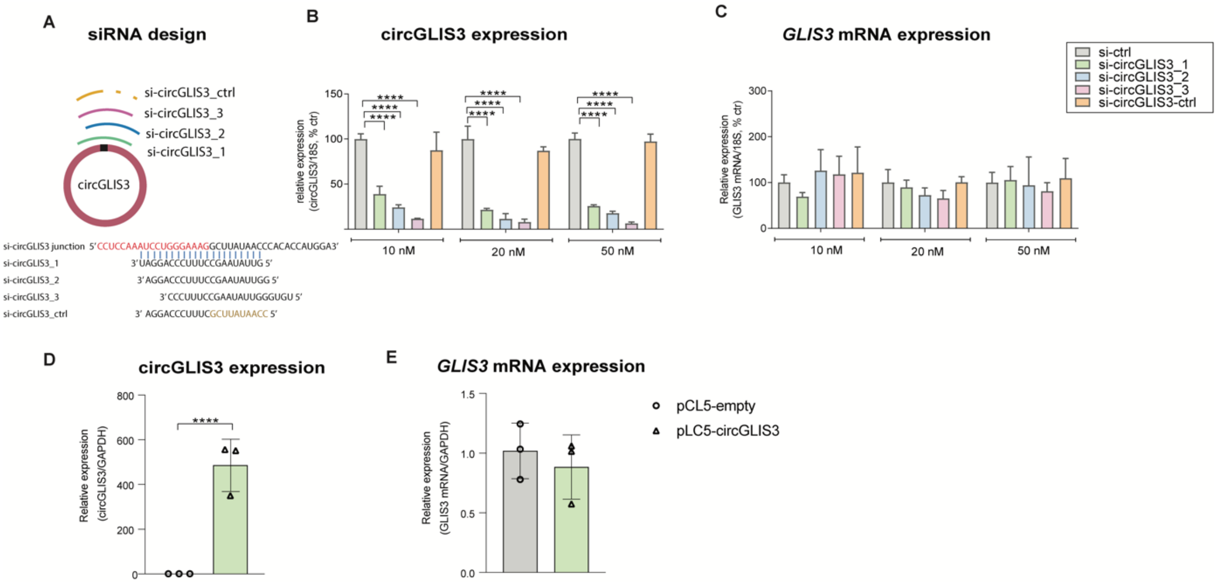


**Figure S2. Modulation of circGLIS3 levels in human dermal fibroblasts.** (**A**) Illustration of siRNA design for circGLIS3 knockdown. We designed three siRNAs targeting circGLIS3 at junction sequence. The red and black sequences show the corresponding 5' and 3' exon sequences forming the junction. A non-targeting siRNA and a siRNA partially targeting the circGLIS3 BSJ (mismatched sequences in the siRNA are shown in brown) were used as controls. qRT-PCR analysis of circGLIS3 (**B**) and *GLIS3* mRNA (**C**) expression in human fibroblasts transfected with 10, 20, or 50 nM of siRNAs targeting circGLIS3 BSJ and the siRNA controls for 24 hours. qRT-PCR analysis of circGLIS3 (**D**) and *GLIS3* mRNA (**E**) expression in human fibroblasts transfected with 250 ng circGLIS3 overexpression plasmid (pCL5-circGLIS3) or empty vector (pCL5-empty) (**F**) for 48 hours. ****P < 0.0001 by two-tailed Student's t-test.

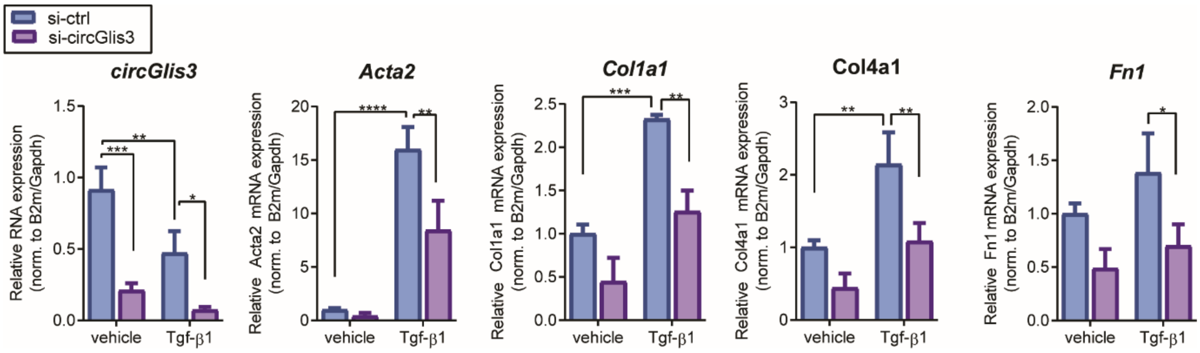


**Figure S3. circGlis3 regulates Tgf-β1 target genes in mouse fibroblasts.** qRT-PCR of circGlis3, *Acta2, Col1a1, Col4a1, and Fn1* mRNA in mouse fibroblasts transfected with si-ctrl or si-circGlis3 for 24 hours and then stimulated with Tgf-β1 (5 ng/ml) for another 24 hours (n=4). *P < 0.05, **P < 0.01, ***P < 0.001, and ****P < 0.0001 by one-way ANOVA and Dunnett's multiple comparisons test.

### SUPPLEMENTAL TABLES

#### Table S1. Human sample information

| Donor | Sex | Age | Ethnicity | Body Location | Experiment |
|---|---|---|---|---|---|
| 1 | F | 66 | Caucasian | Lower leg | RNA sequencing |
| 2 | M | 69 | Caucasian | Lower leg | RNA sequencing |
| 3 | F | 67 | Caucasian | Lower leg | RNA sequencing |
| 4 | M | 69 | Caucasian | Lower leg | RNA sequencing |
| 5 | F | 64 | Caucasian | Lower leg | RNA sequencing |
| 6 | F | 60 | Caucasian | Lower leg | qRT-PCR |
| 7 | F | 66 | Caucasian | Lower leg | qRT-PCR |
| 8 | F | 60 | Caucasian | Lower leg | qRT-PCR |
| 9 | F | 67 | Caucasian | Lower leg | qRT-PCR |
| 10 | F | 65 | Caucasian | Lower leg | qRT-PCR |
| 11 | M | 26 | Caucasian | Lower back | Cell isolation, RNA sequencing |
| 12 | F | 30 | Caucasian | Lower back | Cell isolation, RNA sequencing |
| 13 | M | 45 | Caucasian | Lower back | Cell isolation, RNA sequencing |
| 14 | F | 43 | Caucasian | Lower back | Cell isolation, RNA sequencing |
| 15 | M | 22 | Caucasian | Lower back | Cell isolation, RNA sequencing |
| 16 | F | 48 | Caucasian | Lower back | Cell isolation, qRT-PCR |
| 17 | M | 27 | Caucasian | Lower back | Cell isolation, qRT-PCR |
| 18 | F | 24 | Caucasian | Lower back | Cell isolation, qRT-PCR |
| 19 | M | 24 | Caucasian | Lower back | Cell isolation, qRT-PCR |
| 20 | F | 46 | Caucasian | Lower back | Cell isolation, qRT-PCR |
| 21 | M | 50 | Caucasian | Lower back | LCM, qRT-PCR |
| 22 | M | 28 | Caucasian | Lower back | LCM, qRT-PCR |
| 23 | M | 35 | Caucasian | Lower back | LCM, qRT-PCR |
| 24 | F | 58 | Caucasian | Lower back | LCM, qRT-PCR |
| 25 | F | 53 | Caucasian | Lower back | LCM, qRT-PCR |
| 26 | F | 37 | Caucasian | Lower back | LCM, qRT-PCR |
| 27 | M | 23 | Caucasian | Lower back | LCM, qRT-PCR |
| 28 | F | 48 | Caucasian | Abdomen | *Ex vivo* wound model |
| 29 | F | 56 | Caucasian | Thigh | *Ex vivo* wound model |
| 30 | F | 43 | Caucasian | Abdomen | *Ex vivo* wound model |
| 31 | F | 49 | Caucasian | Abdomen | Fibroblast isolation |
| 32 | F | 51 | Caucasian | Thigh | Fibroblast isolation |
| 33 | F | 28 | Asian | Abdomen | qRT-PCR |
| 34 | F | 41 | Asian | Vulva | qRT-PCR |
| 35 | M | 36 | Asian | Chest | qRT-PCR |
| 36 | M | 35 | Asian | Chest | qRT-PCR |
| 37 | M | 32 | Asian | Chest | qRT-PCR |
| 38 | M | 30 | Asian | Vulva | qRT-PCR |
| 39 | F | 64 | Asian | Chest | qRT-PCR |
| 40 | F | 28 | Asian | Chest | qRT-PCR |

F: Female, M: Male, LCM: Laser Capture Microdissection

**Table S3**. **List of reagents used in this study.**

| Primers/Probes | Sequence/Cat. no/Vendor | Experiment |
|---|---|---|
| circGLIS3 | F: AAAGCAGCAGGAGTTTGGAA<br>R: CTCCTTTCAGGCAAAGTCCA; IDT | PCR, Sanger sequencing, SYBR™ qRT-PCR |
| | F:cgGAATTCTAATACTTTCAGGCTTATAACCCACACCATGGA<br>R:cgGGATCCAGTTGTTCTTACCTTTCCCAGGATTTGGAGGAA<br>F2: GAAGCCGCGATTCCAGGTCA<br>R2: GTGTGGGTTATAAGCCTTTC | circGLIS3 overexpression plasmid construction and verification |
| | F:cgGAATTCTAATACTTTCAGGCTTATAACCCACACCATGGA<br>R:cgGGATCCAGTTGTTCTTACCTTTCCCAGGATTTGGAGGAA<br>F2: CAGAACAACGTGGCTGAGAG<br>R2: TGGTGTGGGTTATAAGCCTTT | circGLIS3-MS2 overexpression plasmid construction and verification |
| *GLIS3* | Hs.PT.58.40536191, IDT | SYBR™ qRT-PCR |
| *ACTA2* | Hs.PT.56a.2542642, IDT | TAQMAN™ qRT-PCR |
| *COL1A1* | Hs.PT.58.15517795, IDT | TAQMAN™ qRT-PCR |
| *COL4A1* | Hs.PT.58.15679435, IDT | TAQMAN™ qRT-PCR |
| *FN1* | Hs.PT.58.40005963, IDT | SYBR™ qRT-PCR |
| *18S* | F: CGGCTACCACATCCAAGGAA<br>R: GCTGTAATTACCGCGGCT<br>probe: TGCTGGCACCAGACTTGCCCTC | SYBR™ /TAQMAN™ qRT-PCR |
| *GAPDH* | F: GGTGTGAACCATGAGAAGTATGA<br>R: GAGTCCTTCCACGATACCAAAG<br>probe: AGATCATCAGCAATGCCTCCTGCA | SYBR™ /TAQMAN™ qRT-PCR |
| *ACTB* | F: GTGGCCGAGGACTTT<br>R: CCTGTAACAACGCAT | SYBR™ qRT-PCR |
| *B2M* | F: AAGTGGGATCGAGACATGTAAG<br>R: GGAGACAGCACTCAAAGTAGAA | SYBR™ qRT-PCR |
| Mouse circGlis3 | F: TCATGGCAGCACCACATACT<br>R: GCCACGCTGATCAATATCCG | PCR, Sanger sequencing, SYBR™ qRT-PCR |
| Mouse *Acta2* | Mm.PT.58.16320644, IDT | SYBR™ qRT-PCR |
| Mouse *Col1a1* | Mm.PT.58.7562513, IDT | SYBR™ qRT-PCR |
| Mouse *Col4a1* | Mm.PT.58.31838522, IDT | SYBR™ qRT-PCR |
| Mouse *Fn1* | Mm.PT.58.28716501, IDT | SYBR™ qRT-PCR |
| Mouse *B2m* | R: CAGTATGTTCGGCTTCCCATTC<br>F: TTCTGGTGCTTGTCTCACTGA | SYBR™ qRT-PCR |
| Mouse *Gapdh* | Mm.PT.39a.1, IDT | SYBR™ qRT-PCR |

| | | | |
|---|---|---|---|
| BA-DapB-1zz | 701021; BaseScope™ negative control probe; ACD, Bio-Techne | | In situ hybridization |
| BA-Hs-PPIB-1zz | 710171; BaseScope™ positive control probe; ACD, Bio-Techne | | In situ hybridization |
| BA-Hs-GLIS3-circRNA-E2 | 827481; BaseScope™ custom probe targeting circGLIS3; ACD, Bio-Techne | | In situ hybridization |
| **siRNAs** | **Sequence/Cat. no/Vendor** | | **Experiment** |
| si-ctrl | (AGG UAG UGU AAU CGC CUU G)TT<br>Non-targeting siRNA control; Eurofins | | circGLIS3 knockdown |
| si-circGLIS3_ctrl | (UCC UGG GAA AGC GAA UAU UGG )TT<br>siRNA control partially targeting circGLIS3 BSJ; Eurofins | | circGLIS3 knockdown |
| si-circGLIS3_01 | (AUC CUG GGA AAG GCU UAU AAC) TT<br>siRNA fully targeting circGLIS3 BSJ; Eurofins | | circGLIS3 knockdown |
| si-circGLIS3_02 | (UCC UGG GAA AGG CUU AUA ACC )TT<br>siRNA fully targeting circGLIS3 BSJ; Eurofins | | circGLIS3 knockdown |
| si-circGLIS3_03 | (GGG AAA GGC UUA UAA CCC ACA )TT<br>siRNA fully targeting circGLIS3 BSJ; Eurofins | | circGLIS3 knockdown |
| si-ctrl | D-001210-03-05<br>siRNA non-targeting control; Dharmacon™ | | Mouse circGlis3 knockdown |
| si-circGlis3 | CUC CAA AUC CUG AGA AAG ACU UAU CdTdT<br>siRNA targeting the mouse circGlis3 BSJ; Dharmacon™ | | Mouse circGlis3 knockdown |
| si-ctrl ON-TARGET™plus | D-001810-03-20; Dharmacon™ | | PCPE-1 knockdown |
| si-PCOLCE ON-TARGET™plus | L-011747-00-0005<br>siRNA targeting PCOLCE; Dharmacon™ | | PCPE-1 knockdown |
| **Plasmids** | **Information/Vendor** | | **Experiment** |
| pLC5-empty | pLC5-ciR empty backbone for circRNA overexpression; Geneseed | | circGLIS3 overexpression |
| pLC5-circGLIS3 | circGLIS3 overexpression plasmid; Geneseed | | circGLIS3 overexpression |
| pLO-circGLIS3 | circGLIS3 overexpression plasmid; Creative Biogene | | circGLIS3 pulldown |
| pLO-circGLIS3-MS2 | circGLIS3 - MS2 hairpins overexpression plasmid; Creative Biogene | | circGLIS3 pulldown |
| MS2-CP | Plasmid overexpressing MS2 capture protein; Creative Biogene | | circGLIS3 pulldown |
| pBV-luc | plasmid #16539, Addgene | | Luciferase reporter assay |
| pSBE4-luc | plasmid #16495, Addgene | | Luciferase reporter assay |
| **Antibodies** | **Cat. no/Vendor** | **Dilution** | **Experiment** |
| Anti-human β-Actin | A3854, Sigma-Aldrich | 1:20000 | Western blot |
| Anti-human α-SMA | MAB1420, R&D Systems | 1:1000<br>1:100 | Western blot<br>Immunofluorescence |
| Anti-human COL1A | sc-59772, Santa Cruz | 1:1000 | Western blot |

| | | | |
|---|---|---|---|
| Anti-human PCPE-1 | sc-73002; Santa Cruz | 1:1000<br>1:100<br>1:25 | Western blot,<br>Immunofluorescence<br>Simple Western |
| Alexa Fluor 555 donkey anti-mouse IgG | A31570, Invitrogen | 1:200 | Immunofluorescence |
| **Cytokines** | **Cat. No./Vendor** | **Concentration** | **Experiment** |
| rh-IL-1α | 11349013; ImmunoTools | 20 ng/ml | qRT-PCR |
| rh-IL-6 | 11340060; ImmunoTools | 50 ng/ml | qRT-PCR |
| rh-IL-8 | 11349080; ImmunoTools | 50 ng/ml | qRT-PCR |
| rh-IL-22 | 11340223; ImmunoTools | 30 ng/ml | qRT-PCR |
| rh-IL-36α | 11340362; ImmunoTools | 100 ng/ml | qRT-PCR |
| rh-BMP-2 | 11343273, ImmunoTools | 100 ng/ml | qRT-PCR |
| rh-TNF-α | 210-TA, R&D Systems | 50 ng/ml | qRT-PCR |
| rh-EGF | 11343406; ImmunoTools | 20 ng/ml | qRT-PCR |
| rh-FGF-2 | 11343625; ImmunoTools | 30 ng/ml | qRT-PCR |
| rh-IGF-1 | 11343316; ImmunoTools | 20 ng/ml | qRT-PCR |
| rh-HB-EGF | 259-HE-050/CF; R&D Systems | 20 ng/ml | qRT-PCR |
| rh-PDGF-AA | 11343683 ImmunoTools | 20 ng/ml | qRT-PCR |
| rh-PDGF-BB | 11343673; ImmunoTools | 20 ng/ml | qRT-PCR |
| rh-TGF-β1 | 240-B, R&D Systems | 20 ng/ml,<br>5 ng/ml | qRT-PCR<br>Luciferase reporter assay, Western blot, Immunofluorescence, RIP, CETSA |
| rh-TGF-β2 | 11344750; ImmunoTools | 10 ng/ml | qRT-PCR |
| rh-TGF-β3 | 11344751; ImmunoTools | 20 ng/ml | qRT-PCR |
| rh-VEGFA | 11343663; ImmunoTools | 20 ng/ml | qRT-PCR |
| **Chemicals** | **Cat. No./Vendor** | **Concentration** | **Experiment** |
| Actinomycin D | A9415, Sigma-Aldrich | 5 μg/ml | qRT-PCR |
| Cycloheximide | C4859, Sigma-Aldrich | 5 μg/ml | Simple Western |
| MG132 | M7449, Sigma-Aldrich | 0.5 μM | Simple Western |
| RNaseR | RNR07250, Lucigen | 1U/1μg RNA | PCR |

F: Forward, R: Reverse, BSJ: Backsplice junction, LCM: Laser Capture Microdissection, IDT: Integrated DNA Technologies; ACD: Advanced Cell Diagnostics
